# PDGFRα-directed fibroblast-homing peptides for targeted delivery of nucleic acid therapeutics

**DOI:** 10.64898/2026.10.19.759357

**Authors:** Shuzhi Cheng, Will Weston, Pierpaolo Ala, Jennifer C Chandler, William J Mason, Heloise Vinette, Wendy E Heywood, Rachael Dickman, Francesco Muntoni, Sara Aguti, Haiyan Zhou

**Author notes:** To whom correspondence should be addressed. Correspondence may also be addressed to Sara Aguti.

## Abstract

Efficient and cell type-specific delivery remains a major barrier to the clinical translation of nucleic acid therapeutics (NATs), particularly for fibroblast-driven diseases in which target cells are poorly accessible *in vivo*. By integrating single-cell transcriptomics with cell-surface protein profiling, we identified platelet-derived growth factor receptor alpha (PDGFRα) as a fibroblast-enriched surface receptor. A panel of PDGFA-derived short peptides was designed and screened across multiple human and mouse cell types, leading to the identification of two fibroblast-homing peptides, Pep3 and Pep5, which displayed rapid and preferential uptake in fibroblasts. PDGFRα knockdown reduced peptide internalisation as measured by live-cell imaging, confirming receptor-mediated uptake. To evaluate their potential as delivery vehicles for NATs, a fibroblast-based cellular model of collagen VI-related congenital muscular dystrophy (COL6-CMD) was used for further validation. Lead peptides were conjugated to a splice-switching phosphorodiamidate morpholino oligomer (PMO) targeting the recurrent *COL6A1* c.930+189C>T variant. Peptide–PMO conjugates retained exon-skipping activity and Pep3-PMO enhanced collagen VI protein deposition in patient-derived fibroblasts following repeated treatment compared to unconjugated PMO. Transcriptomic profiling demonstrated no activation of inflammatory, oncogenic, or PDGFRα receptor-associated signalling pathways following peptide-PMO treatment. Collectively, these findings establish PDGFRα-targeting fibroblast-peptides as a promising platform for selective delivery of NATs to fibroblasts.

## INTRODUCTION

Nucleic acid therapeutics (NATs) are a rapidly expanding class of synthetic oligonucleotides that modulate gene expression, with therapeutic applications across neurological, metabolic, neuromuscular disorders and cancer. Two major subclasses, antisense oligonucleotides (ASOs) and small interfering RNAs (siRNAs), are chemically modified, short single- or double-stranded nucleic acids designed to bind specific RNA targets and downregulate, restore, or modulate gene expression through defined molecular mechanisms (1,2). To date, 13 ASOs and 8 siRNAs have received regulatory approval (3). However, the discrepancy between the limited number of approved therapies and the large number of candidates in clinical development highlights persistent challenges that limit the clinical translation of NATs.

A major barrier to the success of RNA-based therapeutics is efficient delivery to the target organ or cell types. Following systemic administration, unconjugated oligonucleotides predominantly accumulate in the liver and kidneys, resulting in limited biodistribution to extrahepatic and extrarenal sites and increasing the risk of adverse events related to the elevated tissue concentration in these organs (4,5). Consequently, only a small fraction of the administered dose reaches therapeutically relevant target cells, underscoring the urgent need for targeted delivery strategies capable of enhancing oligonucleotide uptake in more desirable organs.

One clinically validated targeting strategy employs N-acetylgalactosamine (GalNAc) ligand, which binds the asialoglycoprotein receptor (ASGPR) highly expressed on hepatocytes (6). GalNAc conjugation enhances hepatocyte uptake of ASOs and siRNAs while reducing distribution to non-parenchymal liver cells. This strategy has enabled the approval of four GalNAc-siRNA drugs (Givosiran (7), Lumasiran (8), Inclisiran (9) and Vutrisiran (10)) and one GalNac-ASO (Eplontersen (11)) establishing ligand-mediated targeting as a clinically validated principle.

Building on this success, efforts have expanded towards extrahepatic delivery using ligands targeting cell-surface receptors. Notable exemples include antibodies and antibody fragments directed against transferrin receptor 1 (TfR1), which have enhanced ASO and siRNA delivery to skeletal muscle and heart and facilitated transport across the blood–brain barrier (BBB) (12–15). Platforms developed by Avidity Biosciences and Dyne Therapeutics are undergoing clinical evaluation for Duchenne muscular dystrophy, myotonic dystrophy type 1 and facioscapulohumeral muscular dystrophy (16,17). Additional preclinical strategies target the glucagon-like peptide-1 receptor (GLP1R) in pancreatic β-cells (18) and sortilin in receptor-expressing tissues, including the spinal cord (19).

In contrast, targeted delivery of oligonucleotides to fibroblasts remains unexplored, despite their involvement in numerous diseases. Fibroblasts are mesenchymal cells responsible for extracellular matrix production, maintenance of tissue structural integrity, and critical signalling interactions.

Dysregulated fibroblast function contributes to a wide range of disorders, including fibrotic diseases (20), epidermolysis bullosa (21) and collagen VI-related congenital muscular dystrophies (COL6-CMD) (22). Although ASOs have demonstrated promising *in vitro* activity in fibroblast-associated disorders, inefficient *in vivo* delivery remains a major translational challenge (23–27).

COL6-CMD exemplifies this limitation. These paediatric neuromuscular disorders result from dominant or recessive variants in *COL6* genes and currently lack disease-modifying treatments. We have previously developed ASO therapies capable of correcting common dominant *COL6* variants in patient-derived fibroblasts (24,25,27,28). Despite robust *in vitro* efficacy, *in vivo* delivery of unconjugated ASOs to muscle interstitial fibroblasts (MIFs), the principal collagen VI-producing cells in skeletal muscle, remains inefficient (1) limiting therapeutic development.

In the present study, we developed a fibroblast-targeting strategy based on short peptides directed against platelet-derived growth factor receptor alpha (PDGFRα), a cell-surface receptor enriched in fibroblasts. We demonstrate that these peptides preferentially internalise into fibroblasts *in vitro* and can be conjugated to exon-skipping ASOs while preserving their molecular activity. To our knowledge, this represents the first fibroblast-directed peptide platform developed for targeted oligonucleotide delivery, offering a promising approach to address delivery limitations in COL6-CMD and potentially other fibroblast-driven disorders.

## RESULTS

### PDGFRα was identified through integration of single-cell transcriptomics and cell-surface protein profiling

To identify a fibroblast-enriched cell-surface receptor suitable for targeted ASO delivery, we integrated publicly available single-cell RNA sequencing (scRNA-seq) data with cell-surface protein profiling analyses across multiple relevant cell types. scRNA-seq data from human skeletal muscle samples were obtained from the Human Cell Landscape database (https://db.cngb.org/HCL/). MIFs were identified based on established marker expression, and genes selectively enriched in this population were extracted for further analysis.

Using Ingenuity Pathway Analysis (IPA), we assessed the subcellular localisation of proteins encoded by 600 genes expressed in MIFs. This analysis identified 64 genes encoding plasma membrane associated proteins. To prioritise candidates with potential for selective targeting, genes exhibiting substantial expression in non-target tissues, including liver, kidney, and muscle fibres, were excluded, yielding a refined list of cell-surface proteins preferentially expressed in MIFs.

These transcriptomic findings were subsequently cross-validated with cell-surface protein profiling data generated from various human cell types. Cell-surface protein analyses were performed on two skin fibroblast lines from healthy controls, three skin fibroblast lines derived from COL6-CMD patients carrying mutations in *COL6A1*, *COL6A2*, or *COL6A3*, human hepatic cell line (HepG2), and muscle-derived cell populations, including myoblasts and MIFs isolated from three healthy donor biopsies. Cell-surface proteins were enriched and analysed by mass spectrometry. All samples yielded high-quality datasets, with the exception of two MIF samples; therefore, subsequent analyses included only one MIF sample.

To stringently identify fibroblast-selective surface receptors, two complementary analytical strategies were applied. In the first approach, proteins commonly detected on the surface of on-target cells (control skin fibroblasts, patient-derived skin fibroblasts, and MIFs) were filtered by removing common proteins detected in off-target cells (myoblasts and HepG2 cells). In the second, more stringent approach, proteins detected in on-target cells were filtered by excluding any protein identified in HepG2 cells or in any of the three myoblast samples, without clustering off-target datasets.

Using the first analytical approach, six proteins were identified as being selectively expressed among on-target fibroblast populations. In contrast, application of the second, more stringent filtering strategy yielded a single candidate, platelet-derived growth factor receptor alpha (PDGFRα), as the only candidate consistently detected across all fibroblast populations analysed and was absent from all proteins identified in hepatocytes and in the three myoblast populations (Fig. 1).

**Figure 1:**
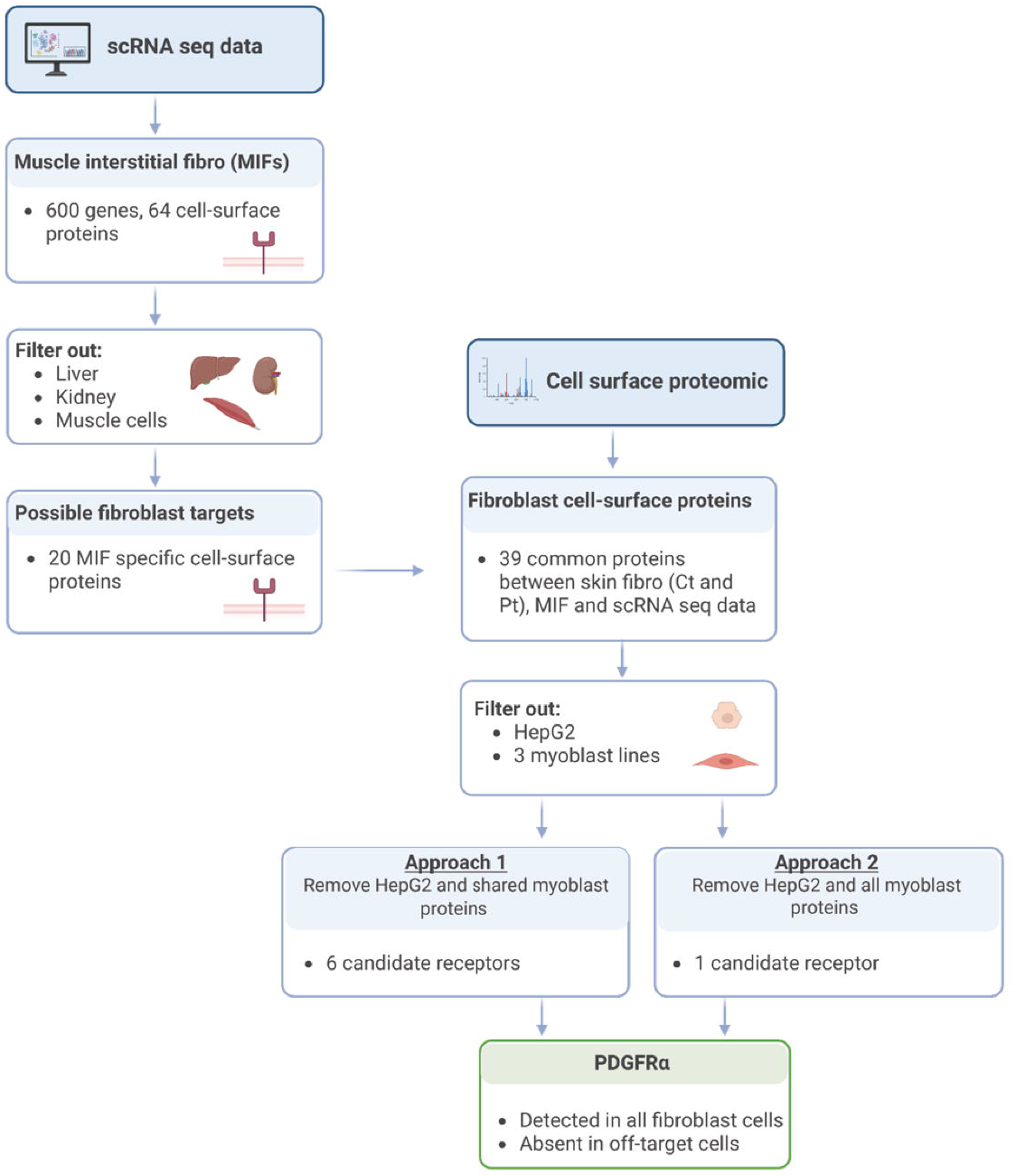
Schematic overview of the workflow used to identify the predominantly expressed receptor in fibroblasts.

This finding is consistent with previous reports describing PDGFRα expression across multiple fibroblast subtypes and supports its selection as a candidate receptor for fibroblast-targeted ASO delivery (37).

### PDGFRα expression was detected across skeletal muscle and skin biopsies and abundantly in fibroblast lines

PDGFRα expression was validated in human skeletal muscle and skin tissues and across a panel of human and mouse cell types to confirm fibroblast-enriched expression consistent with its use as a targeting receptor.

PDGFRα expression in different biopsies and cell lines was measured at both RNA and protein levels, to confirm its enrichment in fibroblast cells. Initially, protein expression of PDGFRα was confirmed in muscle and skin biopsies from healthy controls, and in muscle biopsies from COL6-CMD patients (Fig. 2A-B). As expected, PDGFRα immunostaining was not detected in muscle fibres but rather in the interstitial spaces where fibroblasts reside, as well as in skin biopsies. Furthermore, RNA-level analysis of PDGFRα by qRT-PCR in muscle biopsies from both healthy controls and COL6-CMD patients revealed increased expression in patient samples compared to healthy controls (Fig. 2A).

**Figure 2:**
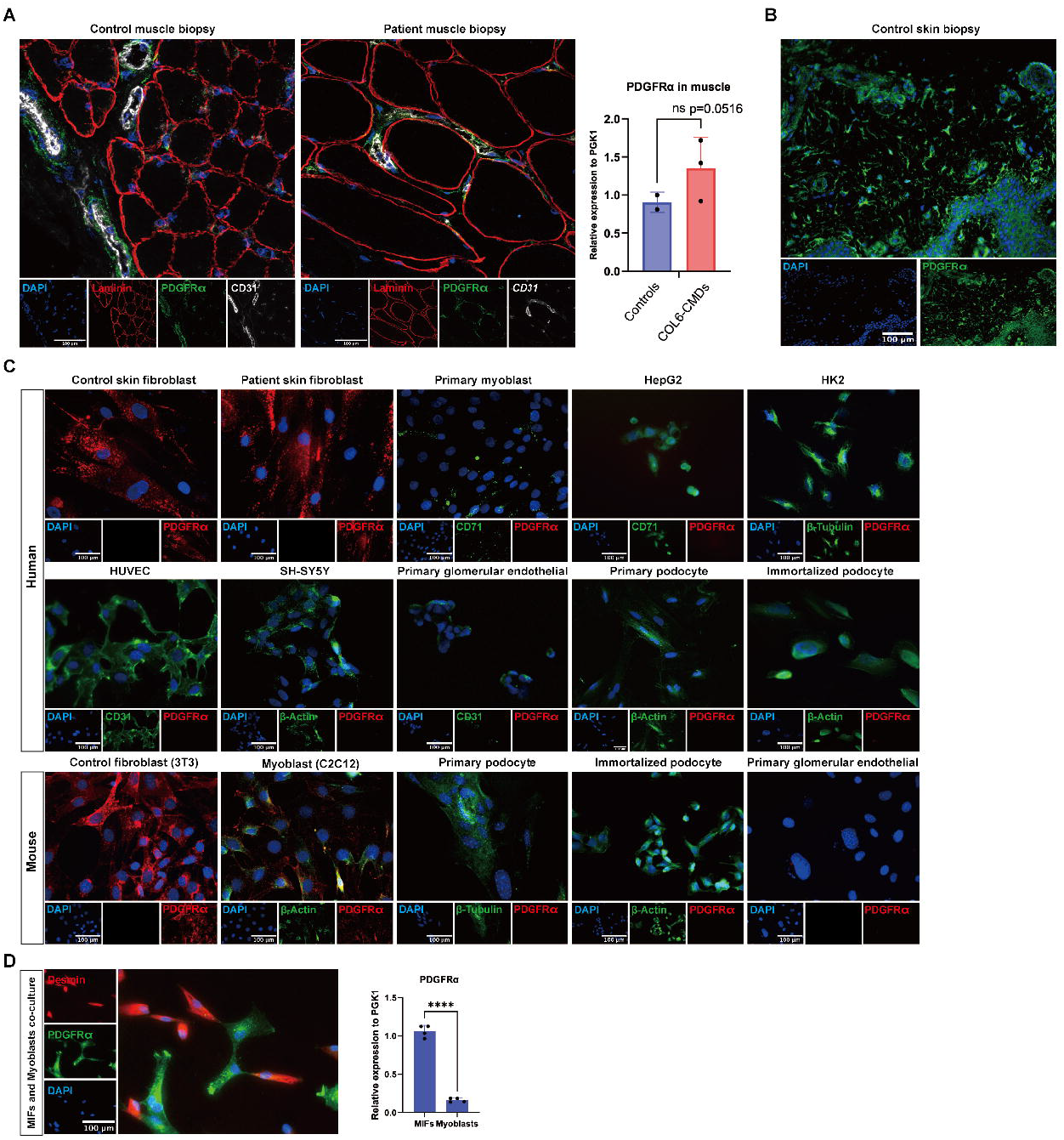
Validation of PDGFRα expression across muscle biopsies and different cell lines: (A) Immunofluorescence staining of PDGFRα (in green), laminin (in red, marker of muscle membrane), CD31 (in grey, marker of endothelium) and nuclei (in blue) in muscle biopsy from non-neuromuscular paediatric donor (control) and COL6-CMD patient. Scale bars: 100 μm. Quantification of PDGFRα expression normalized to PGK1 by RT-qPCR is shown on the right. Control n=2, patients n=3. Data are presented as mean ± SD. Statistical significance was determined using unpaired t-test. (B) Immunofluorescence staining of PDGFRα (in green) and nuclei (in blue) in skin biopsy from non-neuromuscular paediatric donor (control). Scale bars: 100 μm. (C) Immunofluorescence staining of PDGFRα (red) in different human and mouse cell lines. Nuclei were stained with DAPI (blue). Cell markers including CD31, CD71, β-actin and β-tubulin are shown in green. Scale bars: 100 μm. (D) Immunofluorescence staining of PDGFRα (green) and desmin (red) in co-cultured muscle interstitial fibroblasts (MIFs) and myoblasts. Nuclei were stained with DAPI (blue). Scale bars: 100 μm. Quantification of PDGFRα mRNA expression relative to PGK1 measured by RT-qPCR is shown on the right. Data are presented as mean ± SD. Statistical significance was determined using unpaired t-test (**** p < 0.0001).

PDGFRα expression was then evaluated across ten human cell lines, including control and COL6-CMD fibroblasts, myoblasts, HepG2, HUVEC, HK2, immortalized podocytes, primary podocytes, primary glomerular endothelial cells, and SH-SY5Y, and five mouse cell lines, including fibroblasts, myoblasts, immortalized podocytes, primary podocytes, and primary glomerular endothelial cells. Immunostaining revealed PDGFRα protein expression in both human control and patient-derived fibroblasts, as well as in mouse fibroblasts, with lower expression detected in mouse myoblasts. No PDGFRα protein expression was observed in the remaining cell lines (Fig. 2C).

These findings were consistent at the transcript level, with PDGFRα mRNA analysed across all cell lines and showing predominant expression in both human and mouse fibroblasts (Supplementary Fig. S1A). Furthermore, PDGFRα mRNA expression was compared between fibroblasts from healthy controls and COL6-CMD patients carrying mutations in *COL6A1*, *COL6A2*, and *COL6A3*, revealing no significant differences between the groups *in vitro* (Supplementary Fig. S1B).

PDGFRα expression was also assessed in a co-culture system of muscle interstitial fibroblasts (MIFs) and myoblasts derived from three control muscle biopsies. The resulting co-cultures were stained with antibodies against PDGFRα and desmin, a myoblast marker used to distinguish myoblasts from MIFs. Immunostaining showed that PDGFRα expression was restricted to desmin-negative cells, confirming its presence in MIFs but not in myoblasts (Fig. 2D). To evaluate PDGFRα mRNA levels in these co-cultures, MIF- and myoblast-enriched populations were isolated from the co-cultures using conditioned medium (Supplementary Fig. S1C) (35). RT-qPCR analysis revealed significantly higher PDGFRα expression in MIF-enriched cultures compared to myoblast-enriched cultures, supporting the protein-level findings (Fig. 2D).

### Lead PDGFRα ligand peptides were identified across various human and mouse cell lines

Short peptides containing twenty-two amino acids were designed based on the PDGF-A ligand sequence (P04085). Although PDGF-A comprises multiple regions, the cysteine-knot growth factor domain (Cys-Knot) is primarily responsible for receptor recognition (38). Twelve peptides were designed to cover the Cys-Knot region using a “micro-walking” approach, in which each peptide overlaps with the previous one by 12 amino acids (Supplementary Fig. S2, Supplementray table 3). Each peptide was labelled at the N-terminus with the 5-carboxyfluorescein (5-FAM) fluorophore (Eurogentec Ltd) and the uptake of these peptides was assessed in 14 different cell lines: 9 human cell lines (5 primary cells and 4 cell lines) and 5 mouse cell lines (2 primary cells and 3 cell lines). Peptides were incubated at 10 µM concentration for 1h and 3h and measured under fluorescence microscopy (Supplementary Figs. S3 and S4). Among the 12 peptides, Pep3 and Pep5 showed robust uptake in human and mouse fibroblasts with minimal uptake across non-fibroblast cell types (Fig. 3A-C). Uptake was detectable as early as 5 minutes post-incubation (Fig. 3D-E). Based on this data, we designed Pep13 by truncating the lead Pep3 to a shorter 15-amino acid sequence to further evaluate the minimal functional motif. Whilst maintaining its activity of fibroblast-targeting, Pep13 also presents targeting in other cell types, including myoblasts, HepG2 and HK2, therefore was not selected for the subsequent studies.

**Figure 3.**
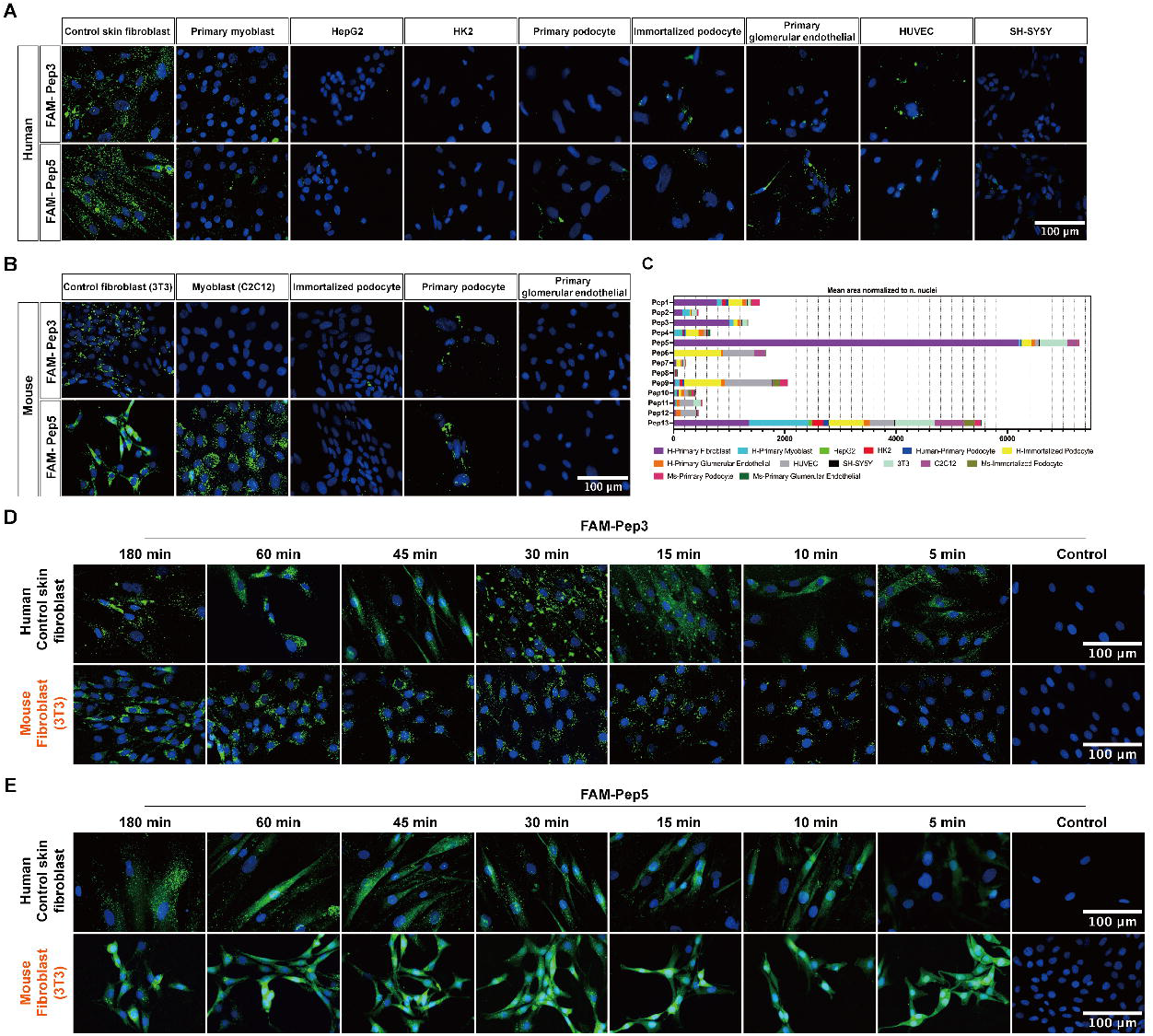
Cellular uptake of candidate peptides across different human and mouse cell lines: (A) Representative fluorescence images showing uptake of FAM-labelled Pep 3 and Pep 5 in human cell lines, including control skin fibroblasts, primary myoblasts, HepG2, HK2, primary podocytes, immortalized podocytes, primary glomerular endothelial cells, HUVEC and SH-SY5Y cells. Nuclei were stained with DAPI (blue). (B) Representative fluorescence images showing uptake of FAM-labelled peptide 3 and peptide 5 in mouse cell types, including control fibroblasts (3T3), myoblasts (C2C12), immortalized podocytes, primary podocytes and primary glomerular endothelial cells. Nuclei were stained with DAPI (blue). (C) Semi-quantitative analysis of fluorescence signal area normalized to the number of nuclei for all tested peptides across various of cell types. (D and E) Representative fluorescence images showing uptake of FAM-labelled Pep3 and Pep5 in human control skin fibroblasts and mouse fibroblasts (3T3) at the indicated time points (5, 10, 15, 30, 45, 60 and 180 min). Untreated cells are shown as controls. Nuclei were stained with DAPI (blue). Scale bars: 100 μm.

### PDGFRα-mediated internalization was confirmed for lead peptides

To assess whether Pep3 and Pep5 uptake is mediated by PDGFRα, we performed PDGFRα siRNA studies to silence its expression in control human fibroblasts. siRNA treatment with 10 nM led to a 70% downregulation of PDGFRα mRNA expression (RT-qPCR) (Supplementary Fig. S5A), with a corresponding reduction in PDGFRα protein staining by 60% (Supplementary Fig. S5B). To confirm the PDGFRα-dependent uptake of Pep3 and Pep5, fibroblasts were treated with PDGFRα siRNA followed by peptides incubation. In addition to Pep3 and Pep5, Pep13 was used as negative control since uptake in a wide range of cell types was observed suggesting a not PDGFRα mediated internalization. As expected, no uptake reduction was observed with Pep13 after PDGFRα knockdown, while reduced Pep3 and Pep5 uptake by ∼70% relative to untreated sample was observed, supporting PDGFRα-dependent uptake for these two lead peptides (Fig. 4A).

**Figure 4.**
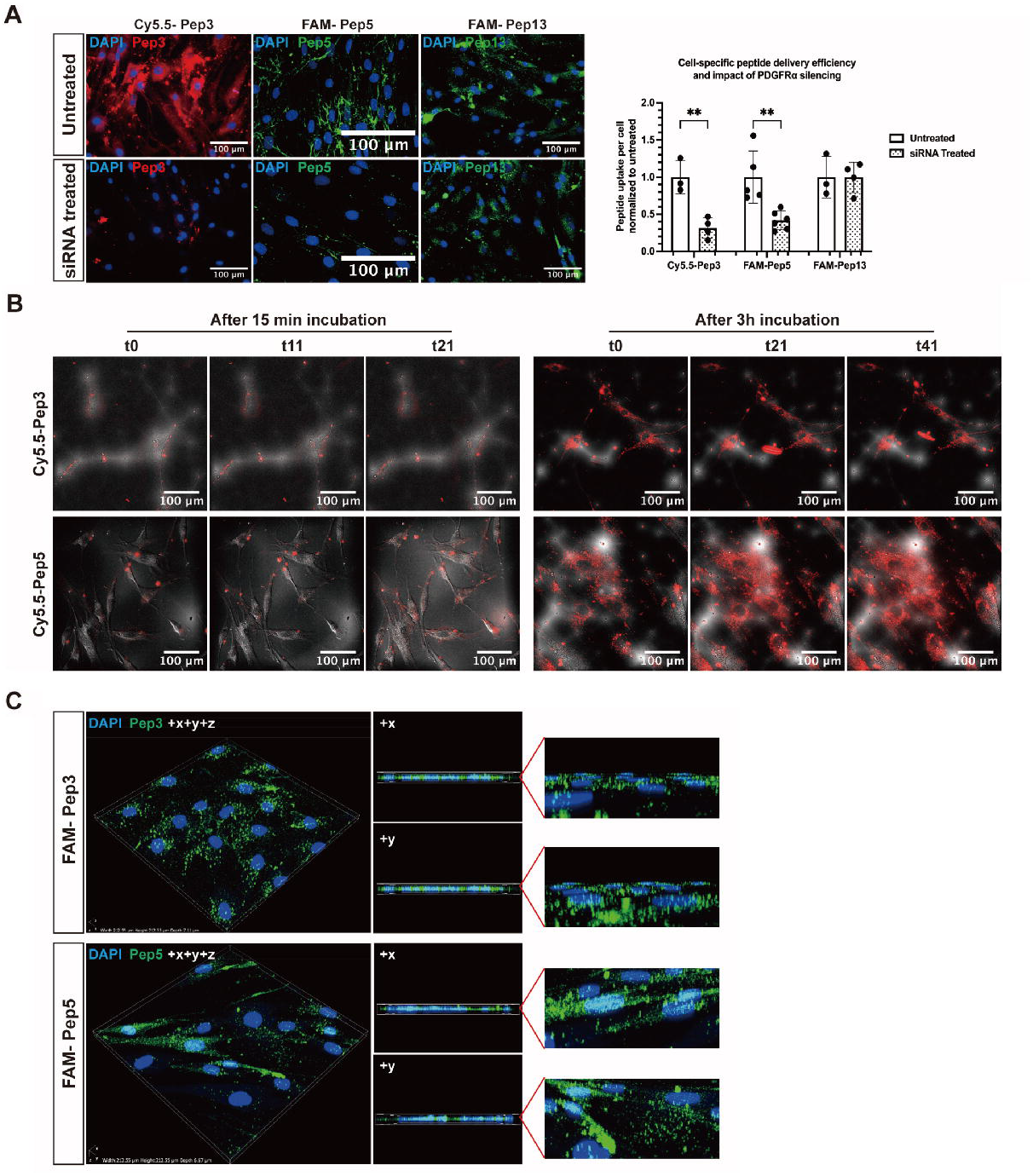
PDGFRα silencing and peptide uptake: (A) Representative fluorescence images showing uptake of Cy5.5-Pep3, FAM-Pep5 and FAM-Pep13 in control fibroblasts under untreated conditions or following PDGFRα siRNA treatment. Nuclei were stained with DAPI (blue). Quantification of peptide uptake per cell normalized to untreated samples is shown on the right. Scale bars, 100 μm. Statistical analysis was performed using Kruskal–Wallis test with Dunn’s multiple comparisons test (**p < 0.01). (B) Representative live-cell fluorescence images of human control fibroblasts following incubation with Cy5.5–Pep3 or Cy5.5–Pep5 (10 μM) for 15 min or 3 h. Time-lapse imaging was performed over 20 min. Cy5.5 fluorescence signal (red) is shown together with corresponding DIC images. (C) Three-dimensional reconstruction of Z-stack confocal images showing cellular distribution of FAM-Pep3 and FAM-Pep5 in fibroblasts. Orthogonal views (+x and +y) are shown to visualize fluorescence signal distribution relative to nuclei (DAPI, blue).

To further verify that receptor binding leads to effective internalization of the peptides, we performed confocal microscopy and time-lapse imaging experiments. Time-lapse imaging indicated initial peptide signal at the plasma membrane within 15 minutes, followed by accumulation in intracellular compartments by 3 hours (Fig. 4B, Supplementary-Videos). Additional confirmation was obtained from Z-stack acquisitions, which revealed that the majority of the fluorescence signal localized to the cytoplasm, with some also detected in the nuclei (Fig. 4C).

### Peptides-ASO conjugation maintained efficiency in correcting altered splicing *in vitro*

To assess that ASO efficacy and uptake are not affected after conjugation with these peptides, we conjugated our lead peptides with exon-skipping ASO that we previously identified to restore the deep intronic variant c.930+189C>T in *COL6A1* gene (24,25). Pep 3 and Pep 5 were conjugated to a phosphorodiamidate morpholino oligomer (PMO9).

The exon-skipping efficiency of unconjugated PMO9 and different conjugates were evaluated in three unrelated patient-derived fibroblast lines carrying the *COL6A1* c.930+189C>T variant under both serum-containing and serum-free gymnotic delivery conditions (Fig. 5A-B). Comparable splicing-correction efficacy, as indicated by the ratio of mutant transcripts to total transcripts, was detected at the RNA level between the unconjugated PMO9 and Pep3- or Pep5-conjugated PMO9, across three patients’ fibroblast lines (Fig. 5A-C).

**Figure 5.**
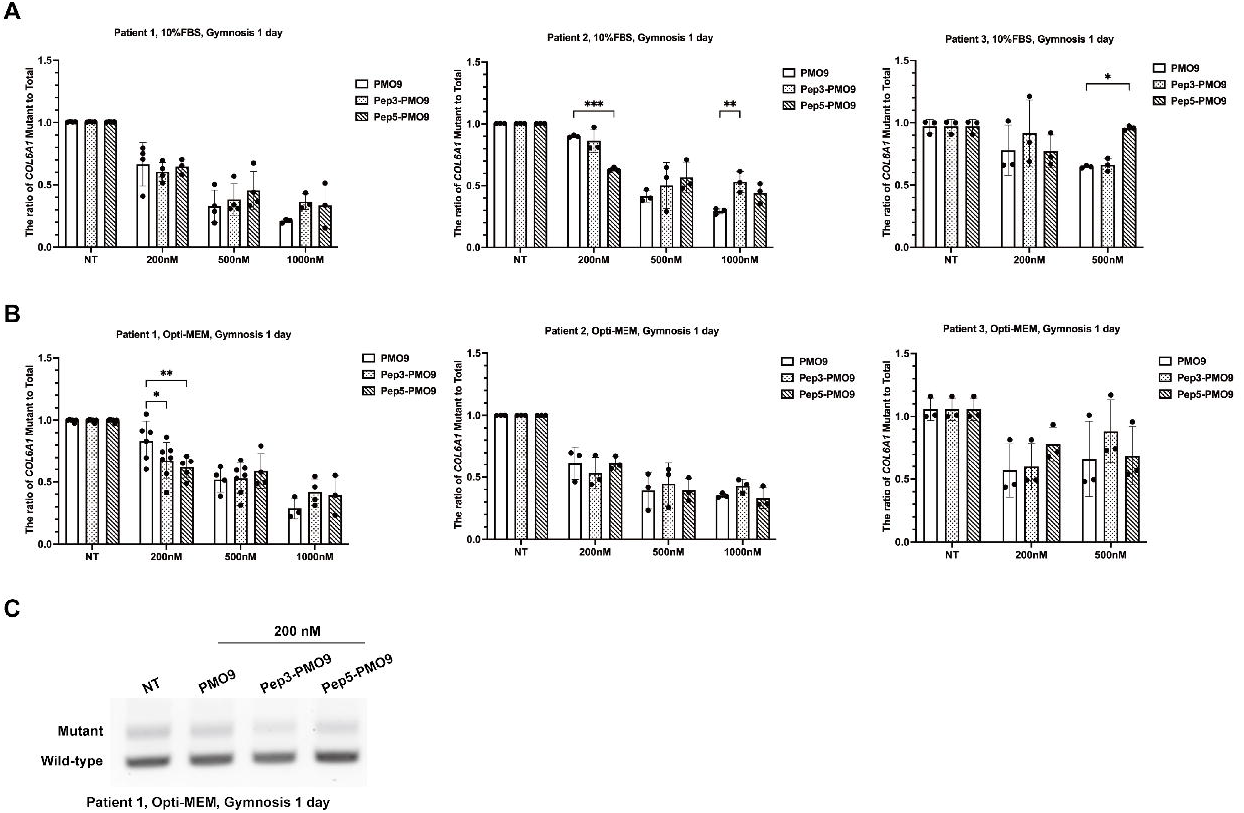
Splice correction by PMO and peptide-PMO conjugates in patient-derived fibroblasts: (A) Quantification of the ratio of mutant to total COL6A1 transcripts in fibroblasts derived from three COL6-CMD patients following gymnotic treatment for 24 hours in medium containing 10% FBS. Cells were treated with PMO9, Pep3-PMO9 or Pep5-PMO9 at the indicated concentrations. NT indicates untreated group. (B) Quantification of the ratio of mutant to total COL6A1 transcripts in fibroblasts derived from three COL6-CMD patients following gymnotic treatment for 24 hours in serum-free medium (Opti-MEM). Cells were treated with PMO9, Pep3-PMO9 or Pep5-PMO9 at the indicated concentrations. NT indicates untreated group. Data are presented as mean ± SD. Statistical significance was determined using one-way ANOVA and post-Bonferroni test (* p<0.05, ** p<0.01,*** p < 0.001).(C) Representative PCR analysis of *COL6A1* transcripts in patient 1 fibroblasts treated with PMO9, Pep3-PMO9 or Pep5-PMO9 at 200 nM under serum-free conditions. Mutant and wild-type bands are indicated.

To investigate whether the transcript-level changes were associated with altered collagen VI deposition, immunofluorescence analysis was performed in one patient’s fibroblasts to assess ECM-associated and total collagen VI distribution following treatment with unconjugated PMO9 or peptide–PMO conjugates (Fig. 6 and Supplementary Fig. S6). Representative fluorescence images showed increased collagen VI deposition following two consecutive treatments (24h + 48h) compared with single treatment (24h), particularly following Pep3–PMO9 treatment (Fig. 6A). Semi-quantitative analysis supported these observations, demonstrating increased ECM-associated collagen VI-positive area and increased total collagen VI-positive area following two consecutive Pep3–PMO9 treatments compared with untreated cells (Fig. 6B). Together, these findings suggest that repeated treatment with Pep3–PMO9 enhances collagen VI deposition in COL6-CMD fibroblasts compared to unconjugated ASO.

**Figure 6.**
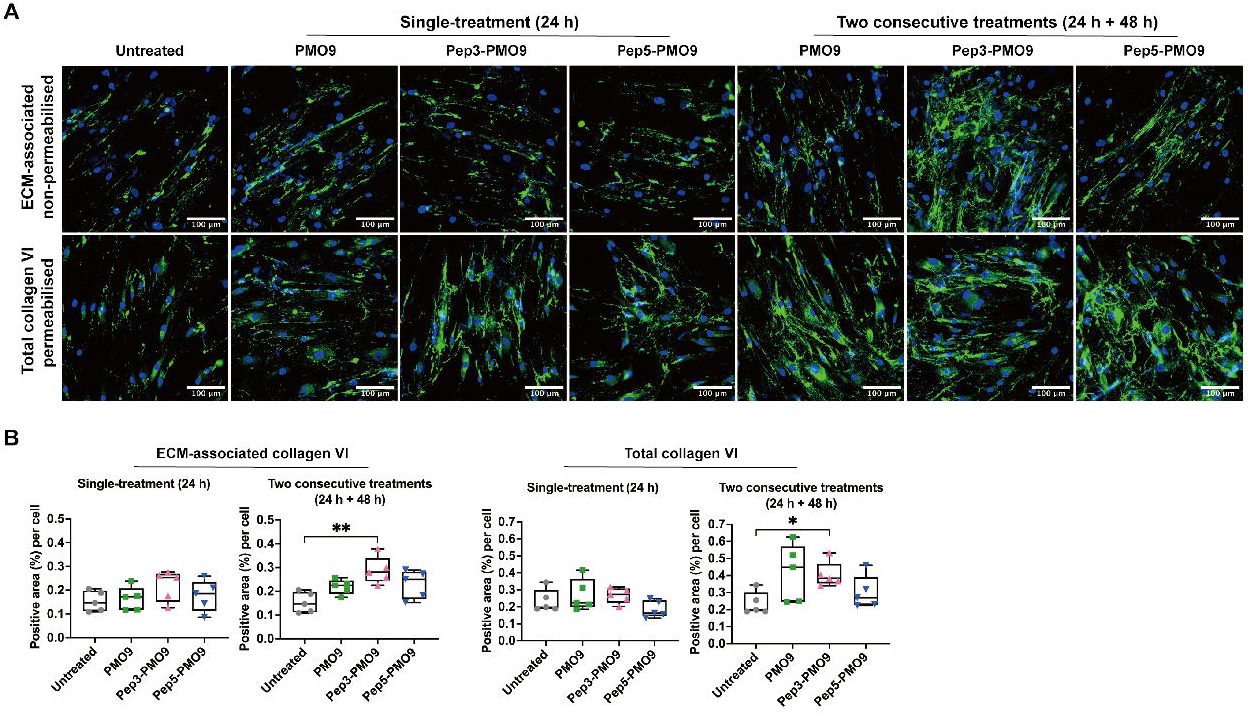
COL6 protein restoration after unconjugated and peptide-PMO conjugates in patient’s fibroblasts: (A) Representative immunofluorescence images showing ECM-associated collagen VI under non-permeabilised conditions and total collagen VI under permeabilised conditions following single treatment (24 h) or two consecutive treatments (24 h + 48 h) with unconjugated PMO9, Pep3-PMO9 or Pep5-PMO9 in COL6-RD fibroblasts. Collagen VI staining is shown in green, and nuclei were counterstained with DAPI (blue). Scale bar, 100 μm. (B) Semi-quantitative analysis of ECM-associated collagen VI and total collagen VI staining following single treatment or two consecutive treatments. Quantification was performed as collagen VI-positive area (%) normalised to cell number. Each point represents an individual image field. Statistical analysis was performed using Kruskal–Wallis test with Dunn’s multiple comparisons test (*P < 0.05, **P < 0.01).

### Peptide–PMO conjugates presented no cytotoxicity *in vitro*

To evaluate whether PMO9 and peptide-PMO conjugates induce cytotoxicity, lactate dehydrogenase (LDH) release assays were performed following gymnotic delivery for 24 hours. HepG2 cells, healthy control fibroblasts and patient-derived fibroblasts were treated with increasing concentrations of PMO9, Pep3-PMO9 and Pep5-PMO9.

No concentration-dependent increase in LDH release was observed for PMO9 or peptide-PMO conjugates in HepG2 cells or fibroblasts (Supplementary Fig. S7), indicating that these compounds do not induce detectable cytotoxicity under the tested conditions. In contrast, treatment with lysis buffer, as positive control, resulted in a strong LDH signal, confirming assay sensitivity.

### Peptide-conjugated PMOs did not activate PDGFRα signalling or altered global transcription

To understand whether the PDGFRα signal pathway is activated by the binding of the short ligand peptides, in addition to the global transcriptomic profiling of ASO correction on mutant transcripts in *COL6A1* gene in patient’s fibroblasts, we performed bulk RNA sequencing in three patients’ fibroblasts lines, treated with different concentrations of Pep-PMO9. Principal component analysis (PCA) showed clear separation of samples according to patient origin, while treated samples remained closely clustered with their respective untreated (NT) controls within each patient group (Fig. 7A), indicating that neither naked PMO9 nor peptide-conjugated PMOs induced any major global transcriptional shift.

**Figure 7.**
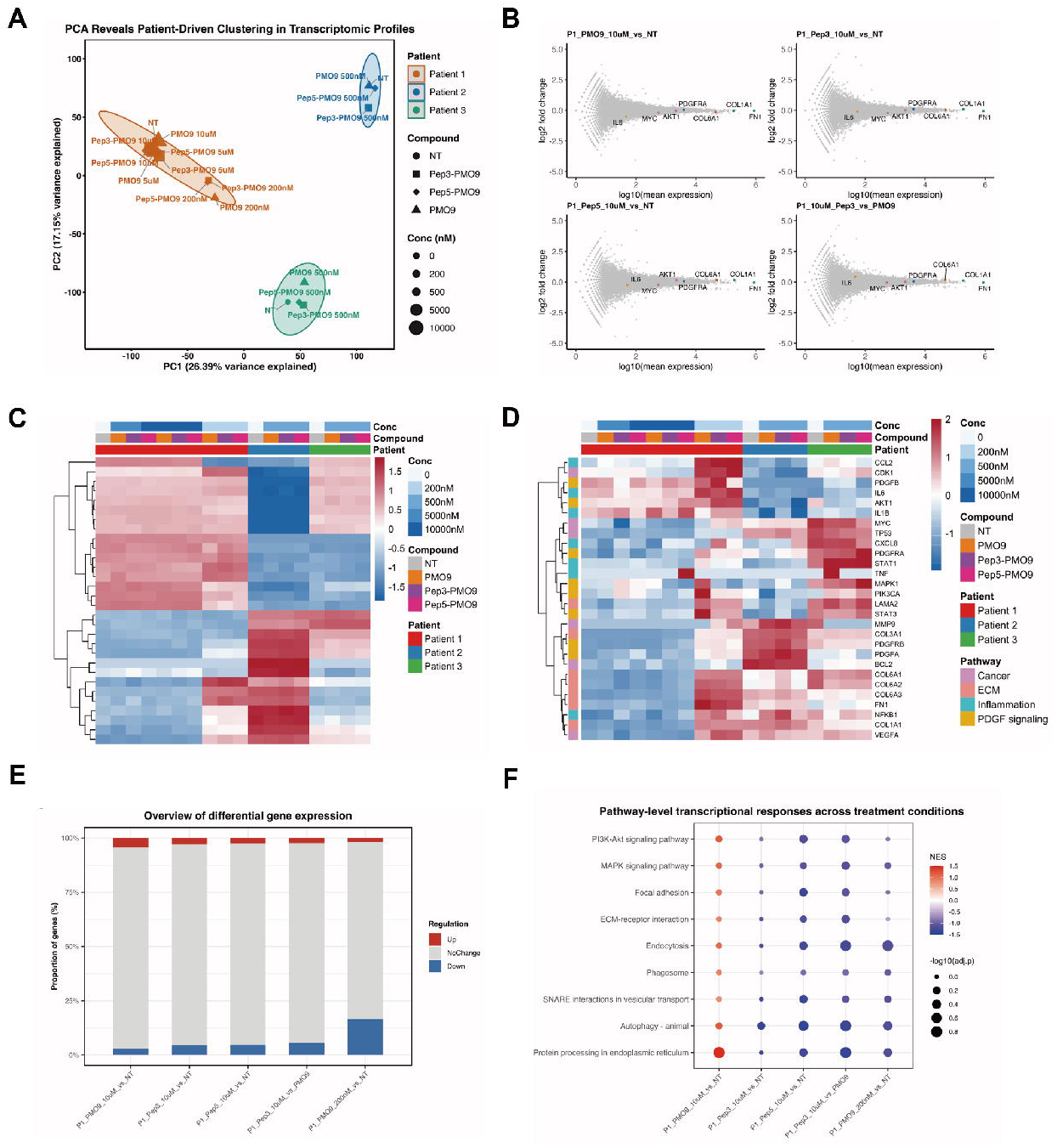
Global transcriptomic, gene expression and pathway-level analysis across treatment conditions: (A) Principal component analysis (PCA) of RNA-seq data from patient-derived fibroblasts treated with PMO9, Pep3-PMO9, and Pep5-PMO9 at indicated concentrations. Samples are coloured by patient and shaped by treatment condition. PC1 and PC2 explain 26.39% and 17.15% of the variance, respectively. (B) MA plots showing gene expression changes for selected comparisons (PMO9 vs NT, Pep3-PMO9 vs NT, Pep5-PMO9 vs NT, and Pep3-PMO9 vs PMO9). Each point represents a gene, plotted as log₂ fold change versus log₁₀ mean expression. Selected genes (*PDGFRA*, *AKT1*, *IL6*, *COL1A1*, *FN1*, *MYC*) are highlighted. (C) Heatmap of the top variable genes across all samples, showing hierarchical clustering based on variance-stabilized (VST) expression values. Samples are annotated by concentration, compound, and patient origin. (D) Heatmap of selected genes associated with PDGFRα signalling, inflammation, extracellular matrix (ECM), and cancer-related pathways. Genes are grouped by functional categories as indicated. (E) Overview of differential gene expression across selected comparisons, showing the proportion of upregulated, downregulated, and unchanged genes. (F) Gene set enrichment analysis (GSEA) showing pathway-level transcriptional responses across treatment conditions. Dot size represents -log10 (adjusted P value), and colour indicates normalized enrichment score (NES).

Consistent with this observation, MA plot analysis across all comparisons showed that the vast majority of genes remained centred around zero log₂ fold change, with no evidence of systematic expression shifts following treatment (Fig. 7B; Supplementary Figs. S8 and S9).

Only a limited number of genes presented modest changes, and these were not consistently shared across patients or treatment conditions. Key genes associated with PDGFRα signalling (PDGFRα, AKT1), inflammation (IL6), extracellular matrix remodelling (COL1A1, FN1), and proliferation (MYC) remained largely unchanged across all groups.

Unsupervised clustering of highly variable genes further confirmed that transcriptomic variation was dominated by inter-patient differences, with no distinct clustering of peptide-conjugated samples observed (Fig. 7C,D; Supplementary Fig. S10).

In line with this, the proportion of differentially expressed genes remained low across different groups, with the majority of genes classified as unchanged (Fig. 7E). Although variation in the relative proportions of up- and down-regulated genes was observed between comparisons, no consistent pattern was associated with peptide-conjugated versus naked PMO treatments.

At the pathway level, gene set enrichment analysis (GSEA) revealed no consistent activation of canonical PDGFRα downstream signalling pathways, including PI3K–Akt and MAPK signalling, across all treatment conditions (Fig. 7F; Supplementary Fig. S11). Pathways associated with inflammation, extracellular matrix remodelling, and oncogenic processes likewise showed no coordinated enrichment. In contrast, pathways involved in intracellular trafficking and protein processing, including endocytosis, phagosome formation, SNARE-mediated vesicular transport, autophagy, and endoplasmic reticulum protein processing, exhibited variable enrichment across different groups. However, these changes were inconsistent in both magnitude and direction, indicating the absence of a reproducible transcriptional signature associated with peptide conjugation.

Collectively, these findings suggest that conjugation of PDGFRα-targeting fibroblast-homing peptides to PMO9 does not trigger ligand-like receptor activation or induce inflammatory, proliferative, or oncogenic transcriptional programs. Instead, peptide-PMO conjugates were associated with transcriptomic profiles that remained largely comparable to those of unconjugated PMO9 and untreated samples, supporting the biological safety and transcriptional neutrality of this delivery strategy.

## DISCUSSION

Targeted ligand-mediated delivery has emerged as one of the most promising approaches for improving the therapeutic index of nucleic acid in target tissue/cell type (1). The clinical success of GalNAc-conjugated oligonucleotides in hepatocytes has demonstrated the transformative potential of receptor-guided delivery platforms (39). More recently, antibody- and Fab-mediated targeting approaches directed against transferrin receptor 1 (TfR1) have shown encouraging results for delivery to skeletal muscle and the central nervous system (14–40). In contrast, selective delivery strategies for fibroblasts remain largely unexplored despite the central contribution of fibroblast dysfunction to numerous disorders, including fibrotic diseases, muscular dystrophies, and connective tissue disorders. To our knowledge, this study describes the first fibroblast-targeting peptide platform developed for receptor-mediated delivery of oligonucleotide therapeutics. More broadly, it establishes a framework for exploiting fibroblast-enriched surface receptors as entry points for cell-selective oligonucleotide delivery, analogous to the role that GalNAc-ASGPR interactions have played in hepatocyte targeting.

In the present study, we developed a fibroblast-enriched targeting strategy based on platelet-derived growth factor receptor alpha (PDGFRα), identified through integration of single-cell transcriptomic and cell-surface protein profiling datasets. Although PDGFRα is widely recognised as a marker of multiple fibroblast populations (37), its potential as a receptor for targeted oligonucleotide delivery has not previously been explored. While PDGFRα is not exclusively restricted to fibroblasts, it is one of the most fibroblast-enriched surface receptors within the cell populations examined in this study, supporting its suitability as a target receptor for fibroblast-selective delivery. Using a rational peptide-design approach based on the PDGF-A ligand, we identified two lead fibroblast-homing peptides, Pep3 and Pep5, that displayed rapid and preferential uptake in both human and mouse fibroblasts.

PDGFRα knockdown markedly reduced peptide internalisation, supporting a receptor-mediated uptake mechanism rather than nonspecific membrane interaction. Live-cell and Z-stack imaging further demonstrated efficient intracellular accumulation following initial membrane association, indicating that receptor engagement results in productive internalisation. Together, these findings establish proof-of-concept for PDGFRα-directed peptide targeting as a strategy for fibroblast-selective cargo delivery.

To evaluate the translational potential of this approach, we have used collagen VI-related congenital muscular dystrophies (COL6-CMD) as a disease model. Over the past decade, several proof-of-concept studies, including work from our group and others, have demonstrated the therapeutic potential of RNA-based strategies for COL6-CMDs. Exon-skipping ASOs, gapmers, and siRNA approaches have successfully restored collagen VI expression at both RNA and protein levels in patient-derived fibroblasts (24,25,27,28,40–42). However, despite these encouraging *in vitro* findings, translation toward *in vivo* application has remained limited, largely because unconjugated oligonucleotides are inefficiently delivered to muscle interstitial fibroblasts (MIFs), the principal source of collagen VI within skeletal muscle (29). Efficient and selective delivery to fibroblast populations therefore represents a major unresolved challenge for the development of nucleic acid therapeutics in COL6-CMDs.

To address this challenge, lead fibroblast-targeting peptides were conjugated to a previously validated splice-switching PMO targeting the recurrent deep intronic *COL6A1* c.930+189C>T variant (24).

Importantly, peptide-PMO conjugates retained exon-skipping activity following conjugation, demonstrating that peptide conjugation did not compromise PMO activity. Interestingly, enhanced functional benefit was detected at the protein level as indicated by collagen VI ECM staining, suggesting the potentially enhanced therapeutic effect of Pep3-PMO9 compared to unconjugated.

The limited enhancement observed in monolayer fibroblast cultures is perhaps not unexpected. In standard cell culture systems, both unconjugated PMOs and peptide-conjugated PMOs are exposed directly to target cells, thereby minimizing the delivery barriers that exist *in vivo*. Consequently, the principal advantage of cell-targeting ligands, namely selective localisation to target cell populations following systemic administration, is likely to be underestimated *in vitro*. The value of the current platform may therefore become more evident in complex multicellular systems and *in vivo* settings, where biodistribution represents a major determinant of therapeutic efficacy. Another factor that may limit functional efficacy is productive intracellular trafficking following receptor-mediated uptake.

Successful delivery of splice-switching ASO requires not only cellular internalisation but also efficient escape from endosomal compartments and subsequent nuclear localisation. In the current study, peptide conjugation was achieved using click chemistry, generating a stable, non-cleavable linkage between the peptide and PMO cargo. Consequently, if the peptide component becomes trapped within endosomal compartments, the conjugated PMO is also likely to remain sequestered, thereby limiting intracellular release and nuclear delivery (44,45). Therefore, is endosomal entrapment will be confirmed future optimisation of the platform may involve incorporation of cleavable linkers, endosomal escape motifs or trafficking-modifying chemistries designed to enhance intracellular bioavailability of the PMO cargo.

An important and often overlooked consideration for receptor-targeted delivery systems is the possibility that receptor engagement itself may elicit unwanted biological responses. This concern is particularly relevant for PDGFRα, a receptor with established roles in multiple biological processes, including proliferation, migration, extracellular matrix remodelling, and fibrosis. Reassuringly, transcriptomic profiling demonstrated that peptide-PMO treatment did not activate canonical PDGFRα downstream pathways, including PI3K-Akt and MAPK signalling, inflammatory or oncogenic transcriptional programs. Collectively, these findings suggest that Pep3 and Pep5 function primarily as delivery ligands rather than receptor agonists, supporting their potential utility as biologically inert targeting moieties.

Several limitations of the present study should be acknowledged. First, transcriptomic analyses were performed primarily in a descriptive manner because of the limited number of biological replicates available from patient-derived fibroblast lines. Nevertheless, the consistency of global transcriptional patterns across independent patient samples supports the overall conclusions. Second, although the current study demonstrates receptor-mediated fibroblast targeting and proof-of-concept functional delivery, evaluated across a panel of cell types *in vitro*; however, comprehensive assessment of tissue specificity and *in vivo* biodistribution following systemic administration remains necessary to define the therapeutic efficacy and potential off-target distribution of this platform. Future studies should therefore focus on optimisation of linker chemistry, dosing regimens, and systemic *in vivo* biodistribution and therapeutic validation in relevant model systems.

In conclusion, we identified PDGFRα as a fibroblast-enriched receptor and developed two PDGFRα-targeting fibroblast homing peptides capable of mediating receptor-dependent cellular uptake of oligonucleotide cargo. Peptide-PMO conjugates retained splice-switching activity, enhanced collagen VI restoration at the protein level under selected conditions, and did not induce detectable cytotoxic, inflammatory or PDGFRα-associated transcriptional responses. These findings establish a proof-of-concept platform for fibroblast-targeted nucleic acid delivery and provide a foundation for future optimisation and *in vivo* evaluation. More broadly, this strategy may enable targeted therapeutic intervention across a wide spectrum of fibroblast-driven genetic, degenerative and fibrotic disorders in which inefficient delivery to fibroblast populations currently limits the translational potential of RNA therapeutics.

## MATERIAL AND METHODS

### Human samples

Written informed consent was obtained from all subjects for the utilization of different primary cell lines, skin biopsies and muscle biopsies samples. This study was conducted in accordance with the Declaration of Helsinki.

Fibroblast lines were established from skin biopsies from paediatric subjects without neuromuscular conditions and patients carrying phatogenic variants in *COL6A1*, *COL6A2* or *COL6A3* genes.

Myoblast lines were established from a muscle biopsy from a paediatric subject without neuromuscular conditions. Established skin fibroblast lines and myoblast lines were supplied by the MRC Centre for Neuromuscular Disease Biobank London (REC reference number: 06/Q0406/33). The use of cells as a model system to study pathogenesis and therapeutic strategies for Neuromuscular Disorders was under etich: REC reference 13/LO/1826.

Primary human podocytes and glomerular endothelial cells were obtained from human fetal kidney tissue provided by the Human Developmental Biology Resource (HDBR, http://hdbr.org) under ethical approval (REC reference 23/LO/0312), and established as previously described (30).

### Cell culture

The established cell lines of human fibroblasts, HepG2, SH-SY5Y, mouse fibroblasts (3T3), and mouse myoblasts (C2C12) were cultured in Dulbecco’s Modified Eagle’s Medium (DMEM; Sigma-Aldrich, USA, #D6429) supplemented with 10% fetal bovine serum (FBS; Sigma Aldrich, MO, USA, #F7524) at 37°C and 5% CO₂. Cells were passaged upon reaching ∼80% confluency.

Human myoblast lines were maintained in complete Skeletal Muscle Cell Growth Medium supplemented with SupplementMix (PromoCell GmbH, Heidelberg, Germany, #C-39365) at 37°C and 5% CO₂. Cells were passaged at 40% confluency. Myotube differentiation was initiated once cells reached 100% confluency by switching to Skeletal Muscle Cell Differentiation Medium with SupplementMix (PromoCell GmbH, Heidelberg, Germany, #C-23061) for 6 days, under standard incubation conditions (37°C, 5% CO₂).

Stable human umbilical vein endothelial cells (HUVECs) were cultured in Endothelial Cell Growth Medium with SupplementMix (PromoCell GmbH, Heidelberg, Germany, #C-22010) at 37°C and 5% CO₂, and were passaged at ∼80% confluency.

Immortalized human proximal tubular epithelial cell lines (HK-2) were maintained in Dulbecco’s Modified Eagle Medium/Ham’s F-12 (DMEM/F-12; Sigma-Aldrich, MO, USA, #D6421) supplemented with 10% FBS at 37 °C and 5% CO₂. Cells were passaged at ∼80% confluency (31).

Conditionally immortalized human podocyte lines were cultured in RPMI 1640 Medium (Thermo Fisher Scientific, Waltham, MA, USA, #21875091), supplemented with 10% FBS and 1% Insulin-Transferrin-Selenium (ITS; 100X; Thermo Fisher Scientific, Waltham, MA, USA, #41400045) at 33°C in 5% CO₂ (32). Upon reaching ∼60% confluency, cells were transferred to 37°C for differentiation over 14 days, with culture medium replaced every two days.

Conditionally immortalized mouse podocyte lines were cultured in RPMI 1640 supplemented with 10% FBS, 1% Penicillin/Streptomycin solution (Gibco, Thermo Fisher Scientific, Paisley, UK, #10378016), and recombinant mouse interferon-γ (IFN-γ; Thermo Fisher Scientific, Waltham, MA, USA, #PMC4033) at 33°C in 5% CO₂ to maintain proliferation. Upon reaching ∼60% confluency, cells were transferred to 37°C and cultured in the same medium without IFN-γ to induce differentiation over 14 days, with culture medium replaced every two days (33).

Primary human podocytes were maintained in RPMI 1640 supplemented with 10% FBS, 1% Penicillin/Streptomycin solution, 100 nM 1,25-dihydroxyvitamin D₃ (Sigma-Aldrich, St. Louis, MO, USA, #D1530), 1 µM all-trans retinoic acid (Sigma-Aldrich, #R2625), and 100 nM dexamethasone (Sigma-Aldrich, #D4902) at 37 °C in 5% CO₂.

Primary mouse podocytes were generated as previously described (34). Cells were maintained in RPMI 1640 supplemented with 10% FBS, 1% penicillin/streptomycin, and 1% ITS at 37 °C in 5% CO₂.

Primary human glomerular endothelial cells and primary mouse glomerular endothelial cells (C57-6014G; Cell Biologics) were maintained using complete endothelial cell culture medium (H1168 for human cells and M1168 for mouse cells; Cell Biologics, Chicago, IL, USA) at 37 °C in 5% CO₂.

Muscle biopsies from three healthy paediatric donors were used to generate co-cultures enriched in myoblasts and MIFs, following the protocol described by Goetsch et al (35).

### Cell-surface protein profiling

Different cell lines were used for cell-surface protein extraction, including fibroblasts from three COL6 patients (harbouring pathogenic variants in *COL6A1*, *COL6A2*, and *COL6A3*), two healthy control fibroblast lines, HepG2, three myoblast-enriched co-cultures, and three fibroblast-enriched co-cultures. Cells were seeded in 15 cm dishes using appropriate growth media to achieve >90% confluency the following day. Cell-surface proteins were extracted according to the manufacturer’s protocol using the Pierce Cell Surface Protein Biotinylation and Isolation Kit (Thermo Fisher Scientific, Waltham, MA, USA, #A44390). Biotinylated proteins were subsequently isolated, digested, and purified using the EasyPep Mini MS Sample Prep Kit (Thermo Fisher Scientific, Waltham, MA, USA, #A40006), following the provided instructions. Peptide concentrations were quantified using the Quantitative Colorimetric Peptide Assay Kit (Thermo Fisher Scientific, Waltham, MA, USA, #A23275), and the samples were analysed using a Synapt G2 mass spectrometer as described previously (36). Ultra-definition mass spectrometry (UDMSE) analysis was performed as previously described (36). Briefly, 0.3 μg of peptides were loaded onto a Waters NanoAquity liquid chromatography (LC) system and separated over 60 minutes on a 75 μm × 150 mm, 1.7 μm Peptide BEH C18 analytical column (Waters, UK), then injected into a Synapt-G2-Si mass spectrometer with ion mobility separation (IMS) (Waters, UK).

Raw data were imported to ProteinLynx (Waters, UK). Processed spectra were searched against the most recent published UniProt human reference proteome database with manual addition of porcine trypsin (P00761). Enzyme digestion was set to trypsin, maximum missed cleavage number was set to 3. Cysteine carbamidomethylation was set as a fixed modification and methionine oxidation, set as a variable modifications. The false discovery rate (FDR) cut-off was set at 1% at both peptide and protein level. Protein quantification was performed using unique peptides only. The resulting protein identifications and quantitative data were exported to Excel (Microsoft) for further analysis.

### Peptide synthesis

For the initial peptide screening in human and mouse cell lines, 1 mg of each peptide was synthesized by Eurogentec Ltd with a purity >70% and labelled at the N-terminus with the 5-carboxyfluorescein (5-FAM) fluorophore. Lead peptides were subsequently synthesized by PeptideSynthetics at >95% purity and labelled at the N-terminus with the Cyanine5.5 (Cy5.5).

### Peptide incubation and cell imaging

Cells were seeded into 8-well glass chamber slides (Thermo Fisher Scientific) with appropriate growth media at a density designed to reach 80% confluency after 24 hours. The following day, the medium was replaced with 400 µL of fresh media containing 10 µM of each peptide. Cells were incubated at 37°C with 5% CO₂ for various time points, ranging from 5 minutes to 3 hours. After incubation, the medium was removed, and cells were washed with PBS and fixed with 4% paraformaldehyde (PFA; Thermo Fisher Scientific, Waltham, MA, USA, #11586711) for 10 minutes at room temperature (RT). Samples were then stained with 4’,6-diamidino-2-phenylindole (DAPI; Sigma-Aldrich, Gillingham, UK, #D8417) and mounted using Anti-Fade Fluorescence Mounting Medium (Abcam, Cambridge, UK, #ab104135).

Cells were visualized using a Leica DMR fluorescence microscope with MetaMorph imaging software. Z-stack confocal images for three-dimensional reconstruction were acquired using a Zeiss LSM710 confocal microscope with Zen Black software (Zeiss). Time-lapse imaging was performed using the ECLIPSE Ti2 inverted microscope (Nikon) and NIS-Elements Imaging Software (Nikon). The time-lapse experiment was conducted at 15 minutes and 3 hours post-incubation and lasted 20 minutes, with images captured every 30 seconds.

### Semi-quantification of fluorescence signal

Semi-quantitative analysis of fluorescence images was performed using ImageJ version 1.52p (https://imagej.nih.gov/ij/). Images were acquired in 16-bit TIFF format and background subtraction was applied before analysis. Two parameters were quantified: mean fluorescence intensity (arbitrary units, AU) and signal area (pixels). For each condition, at least five fields of view were analysed. The total fluorescence intensity and signal area were normalized to the number of nuclei to obtain mean fluorescence intensity per cell and mean signal area per cell.

### Immunofluorescence staining in cultured cells and tissues

Immunofluorescence staining was performed on cells seeded in 8-well glass chamber slides or in 10 µM muscle and skin cryosections. Following fixation with 4% PFA for 10 minutes at RT, samples were washed with PBS and incubated for 1 hour in blocking buffer (5% goat serum in PBS containing 0.1% Triton X-100). Primary antibodies were diluted in blocking buffer and applied to the samples for 1 hour at RT. The following primary antibodies were used: PDGF Receptor α (D1E1E) XP Rabbit mAb (Cell Signaling Technology, mAb3174; 1:200); Mouse anti-Human CD71 Monoclonal Antibody (Bio-Rad, MCA1148; 1:100); Mouse anti-Human CD31 Monoclonal Antibody (Bio-Rad, MCA1738; 1:100); β-Tubulin Antibody 2.1 (Santa Cruz Biotechnology, sc58886; 1:100); Mouse β-actin Antibody (GeneTex, GTX26276; 1:1000); Desmin Clone D33 (clone D33; Dako, M0760; 1:100); Anti-Laminin ɣ1 Antobody (clone A5; Sigma-Aldrich, MAB1914P, 1:50); Collagen VI α3 (Merck Millipore, MAB1944; 1:2000).

After incubation, samples were washed three times with PBS and then incubated for 1 hour at RT with secondary antibodies: goat anti-rabbit Alexa Fluor 594 (Thermo Fisher Scientific, A11012; 1:500), goat anti-rat Alexa Fluor 594 (Thermo Fisher Scientific, A11007; 1:500) or goat anti-mouse Alexa Fluor 488 (Thermo Fisher Scientific, A11029; 1:500). Following three additional PBS washes, nuclei were stained with DAPI and mounted using Anti-Fade Fluorescence Mounting Medium.

### ASO-peptide conjugation

Peptides 3 and 5 were conjugated with the lead exon-skipping phosphorodiamidate morpholino (PMO-9) oligo designed to correct the pre-mRNA splicing of the deep intronic variant in *COL6A1* (24). The conjugation was achieved via copper-free click chemistry, linking the N-terminal azide of each peptide to the 3’-end cyclooctyne of the PMO. PMO-peptide conjugation was performed by Biosynth Ltd.

### Peptide-ASO conjugate gymnotic delivery and cytotoxicity assay

Three unrelated patient’s fibroblast lines carrying c.930+189C>T variant in *COL6A1* were seeded in 24-well plates in DMEM supplemented with 10% FBS at a density designed to reach approximately 80% confluency after 24 hours. The following day, cells were treated with the indicated concentrations of ASO or peptide–ASO conjugates by gymnotic delivery in 500 µL of same medium or Opti-MEM I Reduced Serum Medium (Life Technologies, Paisley, UK, #31985-062). After 24 hours of incubation, cell pellets were collected for RNA extraction, and culture supernatants were collected for cytotoxicity analysis. Cytotoxicity was assessed using the CyQUANT LDH Cytotoxicity Assay (Thermo Fisher Scientific, Waltham, MA, USA, #C20300) according to the manufacturer’s instructions.

### Immunofluorescence-based functional assay of peptide–ASO conjugates

Fibroblast lines carrying the c.930+189C>T variant in *COL6A1* were seeded in 96-well glass-bottom Sensoplates (Greiner) and cultured to achieve approximately 60% confluency prior to treatment. On the day of treatment, cells were treated with ASO or peptide–ASO conjugates under gymnotic conditions using Opti-MEM for 24 h.

For single-treatment conditions, cells were subsequently maintained in complete medium (DMEM supplemented with 10% FBS) containing L-ascorbic acid (50 μg/mL, Fisher Scientific Ltd, UK; Cat# 17997684) for an additional 48 h prior to analysis. For two consecutive treatment conditions, a second dose of ASO or peptide–ASO conjugates was applied in complete medium containing L-ascorbic acid for a further 48 h.

To assess extracellular collagen VI deposition, cells were fixed and stained under non-permeabilised conditions. To assess total collagen VI (intracellular and extracellular), cells were permeabilised using 0.1% Triton X-100 prior to staining. Immunofluorescence staining was performed as described above.

### PDGFRα siRNA transfection

To downregulate PDGFRα, siRNAs targeting PDGFRα (hs.Ri.PDGFRA.13.1, hs.Ri.PDGFRA.13.2 and; Integrated DNA Technologies, IDT) were used. The siRNA sequences are provided in Supplementary Table 1. Healthy control fibroblasts were seeded in 24-well plates at a density of 5 × 10^4^ cells per well to achieve approximately 80% confluence after 24 hours. Cells were then transfected in Opti-MEM I Reduced Serum Medium using Lipofectamine 2000 (Invitrogen^TM^, UK, #11668027) for 24 hours following the manufacturer’s protocol.

### RNA extraction, reverse transcription, RT-qPCR

Total RNA was extracted from cells or tissues using the RNeasy Mini Kit (Qiagen, Hilden, Germany, #74106) following the manufacturer’s instructions. RNA quality and concentration were assessed using a DS-11 Series Spectrophotometer/Fluorometer (DeNovix). First-strand cDNA synthesis was performed using the High-Capacity RNA-to-cDNA Kit (Applied Biosystems, CA, USA, # 4387406), according to the manufacturer’s protocol. Quantitative real-time PCR (qRT-PCR) was carried out using either Takyon ROX Probe or Takyon ROX SYBR MasterMix (Eurogentec, Liège, Belgium, #UF-RPMT-C0701 and #UF-RSMT-B0701) with 20 ng of cDNA per reaction. For *COL6A1* detection, primers and probes were described in our previous study (24,25). Primer sequences used for SYBR Green based qRT-PCR are listed in Supplementary Table 2.

qRT-PCR was performed on a StepOne Real-Time PCR System (Applied Biosystems). For probe-based assays, the thermal cycling conditions were: 50 °C for 2 min, 95 °C for 10 min, followed by 40 cycles of 95 °C for 15 s and 60 °C for 1 min. For SYBR Green assays, the program consisted of 95 °C for 10 min followed by 40 cycles of 95 °C for 15 s and 60 °C for 1 min, with a final melt curve stage (95 °C for 15 s, 60 °C for 1 min, and 95 °C for 15 s). The relative quantification was measured by ΔΔCt method.

### RNA sequencing and transcriptomic analysis

Patient-derived dermal fibroblasts from three unrelated individuals carrying the heterozygous deep intronic variant c.930+189C>T in *COL6A1* were treated with naked PMO9 or peptide-conjugated compounds (Pep3-PMO9 and Pep5-PMO9) at the indicated concentrations using gymnotic delivery for 1 day, or untreated. Total RNA was extracted as described above and submitted to a commercial provider (Novogene) for library preparation and sequencing on an Illumina platform. Reads were aligned to the human reference genome (hg38/GRCh38), and gene-level count tables and FPKM values were generated.

Downstream analysis was performed in R (v4.3.2). Lowly expressed genes were filtered, and count data were normalized using DESeq2 (v1.42.1). Variance stabilizing transformation (VST) was applied for downstream visualization. Principal component analysis (PCA) was performed to assess global transcriptional variation across samples. Gene expression changes between conditions were visualized using MA plots. For gene level visualization, selected genes representing key biological processes, including disease-related extracellular matrix components (COL6A1, COL1A1, FN1), PDGFRα targeting (PDGFRA), inflammation (IL6), and proliferation-associated signalling (MYC, AKT1), were annotated and highlighted in MA plots.

Given the limited sample size, transcriptomic analyses were primarily interpreted in a descriptive manner, focusing on global expression patterns and pathway-level responses rather than statistical significance. Gene set enrichment analysis (GSEA) was performed using clusterProfiler (v4.10.1) based on ranked log2 fold changes. Pathway analysis focused on predefined pathways of interest, including PDGFRα downstream signalling (PI3K–Akt and MAPK pathways), extracellular matrix and adhesion pathways (ECM–receptor interaction, focal adhesion), and intracellular trafficking and stress-related pathways (endocytosis, phagosome, SNARE-mediated vesicular transport, autophagy, and protein processing in the endoplasmic reticulum).

All processed data are provided in the supplementary materials.

### Statistics

GraphPad Prism version 10.0 was used for statistical analysis and graphs design. RT-qPCR data were analysed using one-way ANOVA and post-Bonferroni test or unpaired t-test and presented as mean ± standard deviation (SD). Immunofluorescence staining data were analysed using Kruskal-Wallis test with Dunn’s multiple comparisons test. Differences were considered to be statistically significant at *p ≤0.05; **p ≤ 0.01; ***p ≤ 0.001.

## DATA AVAILABILITY

The data underlying this article will be shared on reasonable request to the corresponding author. The RNA-seq data generated in this study have been deposited in the NCBI Gene Expression Omnibus (GEO) under accession number GSE346574.

## ACKNOWLEDGEMENTS

This work was supported by Muscular Dystrophy UK (19GROPG36-0293 and 22GRO-PG36-0552), LifeArc, UK Research and Innovation (UKRI) Medical Research Council (MR/X008029/1 and MR/Y008405/1), Rosetrees Trust (Seedcorn2023-100173), UCL Technology Fund (UTF-23-001) and Million Dollar Bike Ride Grant Program (MDBR-24-23-CMD) and the National Institutes of Health Research Biomedical Research Centre at Great Ormond Street Hospital for Children NHS Foundation Trust and University College London. The MRC Centre for Neuromuscular Diseases Biobank, supported by the National Institute for Health Research Biomedical Research Centres at Great Ormond Street Hospital for Children NHS Foundation Trust and at University College London Hospitals NHS Foundation Trust and University College London, is acknowledged for providing patients’ fibroblast cell lines. The authors would like to acknowledge Prof. Deborah Eastwood and Dr. Davis Ferguson for providing heathy control skin and muscle biopsies. The authors would like to acknowledge Dr. Carsten Bonnemann, Dr. Veronique Bolduc and Mr. Fady Guirguis for their scientific advice.

## AUTHOR CONTRIBUTIONS

Haiyan Zhou: Conceptualization, methodology, writing-review & editing. Sara Aguti: Conceptualization, methodology, Investigation (receptor identification and peptide sequences design), writing-original draft. Francesco Muntoni: Conceptualization, methodology, writing-review & editing. Shuzhi Cheng: Conceptualization, methodology, Investigation (conducted receptor validation, peptide screening, peptide internalization, and functional experiments), formal analysis, visualization, writing-original draft. Pierpaolo Ala: Resources (established primary fibroblast and myoblast lines from skin and muscle biopsies). Jennifere C Chandler: Resources (established primary podocytes and glomerular endothelial cell lines from mouse glomeruli). William J Mason: Resources (established primary podocytes and glomerular endothelial cell line from HDBR foetal kidneys). Heloise Vinette: Investigation (performed mass spectrometry analysis). Wendy Heywood: Investigation (performed mass spectrometry analysis).

