## Supplemental information for "PDGFRα-directed fibroblast-homing peptides for targeted delivery of nucleic acid therapeutics"

**Supplementary Table 1:** siRNAs sequences targeting PDGFRα and PDGFRβ.

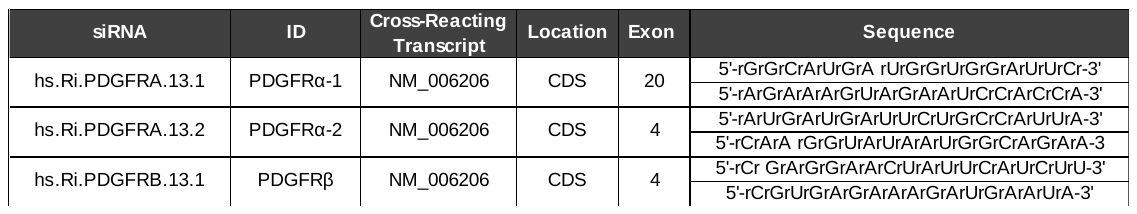

**Supplementary Table 2:** SYBR qRT-PCR primers

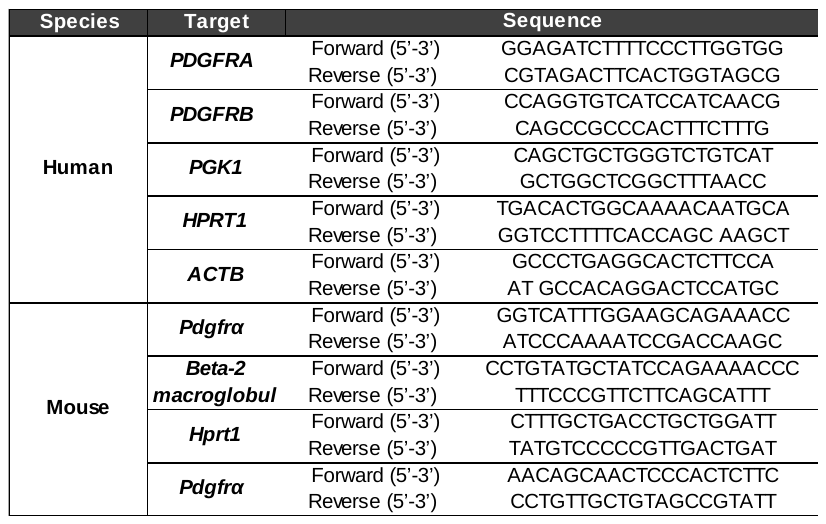

**Supplementary Table 3:** Peptide sequences

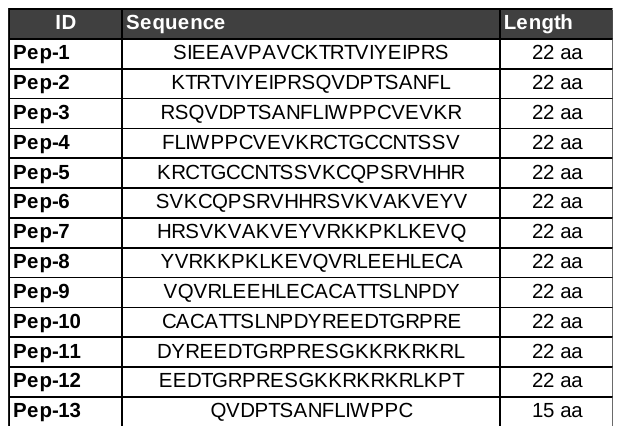

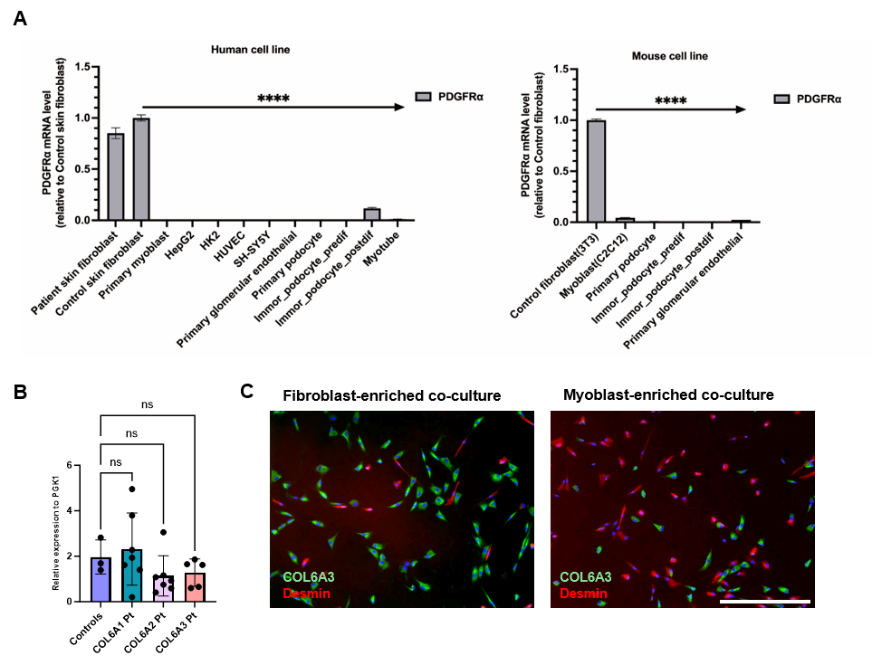

**Supplementary Figure S1:** **PDGFRα expression**.

(A) Relative PDGFRα mRNA expression levels were assessed by RT-qPCR in a panel of human (left) and mouse (right) cell lines. Expression levels are shown relative to fibroblast controls. Data are presented as mean ± SD. Statistical significance was determined using one-way ANOVA and post-Bonferroni test (****P < 0.0001). (B) PDGFRα expression in control skin fibroblasts and in COL6-CMD patients carrying *COL6A1*, *A2* and *A3* mutations. Data are presented as mean ± SD. Statistical significance was determined using one-way ANOVA and post-Bonferroni test. (C) The relative proportions of myoblasts and fibroblasts in fibroblast-enriched and myoblast-enriched co-cultures were quantified by immunostaining for desmin, a myoblast marker, and COL6A3, expressed in fibroblasts but not in myoblasts. Fibroblast-enriched co-cultures contained 83% COL6A3-positive cells and 17% desmin-positive cells, whereas, myoblast-enriched co-cultures contained 65% desmin-positive cells and 35% COL6A3-positive cells. Nuclei were stained with DAPI (blue). Scale bars: 100 μm

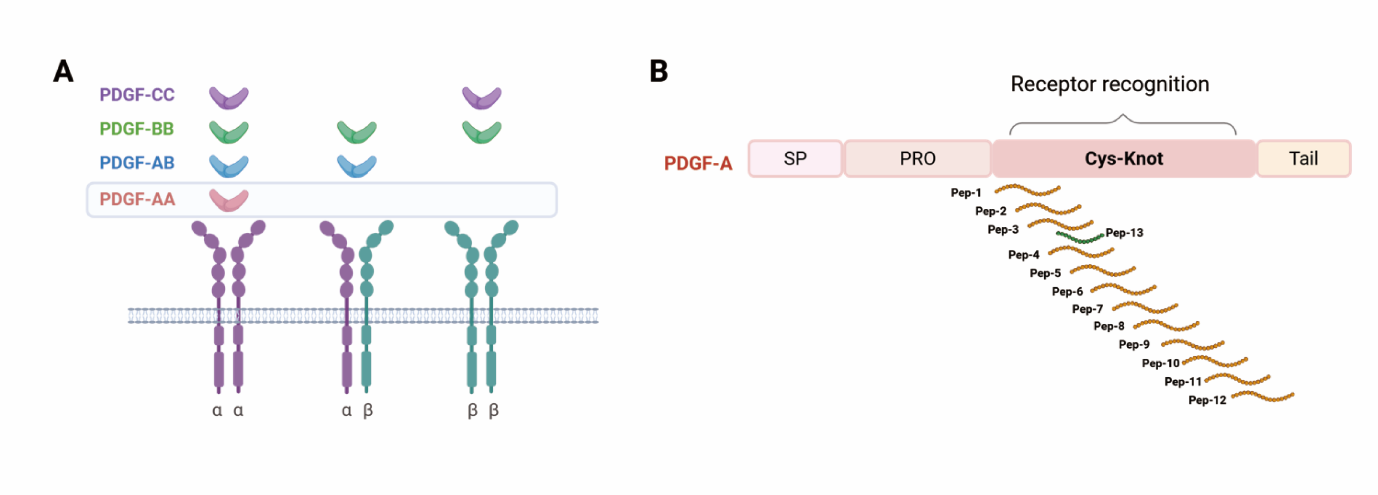

**Supplementary Figure S2: Peptides design strategy.**

(A) Schematic representation of PDGF receptor dimers and their ligand-binding specificity. PDGF-CC, PDGF-BB and PDGF-AB interact with multiple PDGFR dimer combinations, whereas PDGF-AA shows selective binding to the PDGFRαα homodimer. (B) Schematic overview of the peptide-design strategy. Overlapping peptides were designed using a walking approach to span the PDGF-A receptor-binding region within the cysteine-knot growth factor domain.

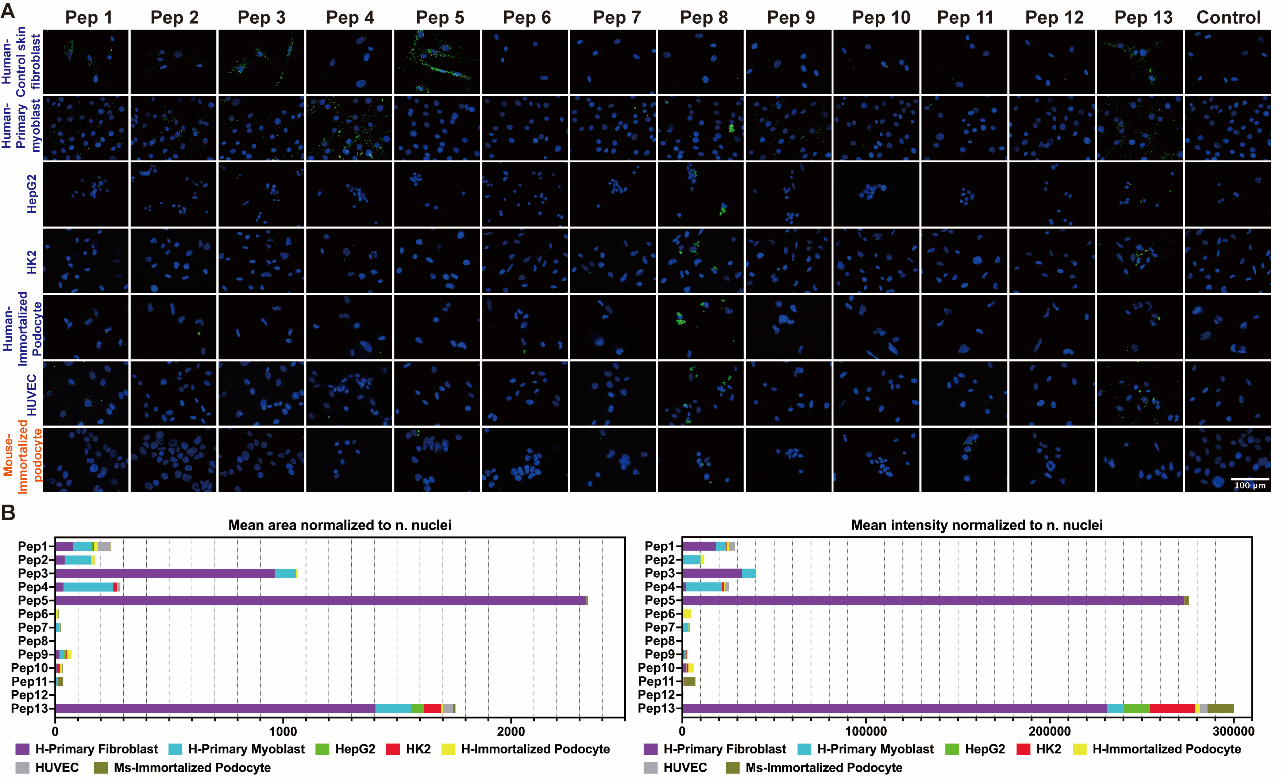

**Supplementary Figure S3: Cellular uptake of peptides across different human and mouse cell lines after 1 hour.**

(A) Representative fluorescence images showing uptake of FAM-labeled peptides in human cell lines, including control skin fibroblasts, primary myoblasts, HepG2, HK2, immortalized podocytes, HUVEC, and also in mouse immortalized podocytes. Nuclei were stained with DAPI (blue). Scale bars, 100 μm. (B) Semi-quantitative analysis of fluorescence signal area (left) and (**C**) intensity (right) normalized to the number of nuclei for all tested peptides across various of cell types.

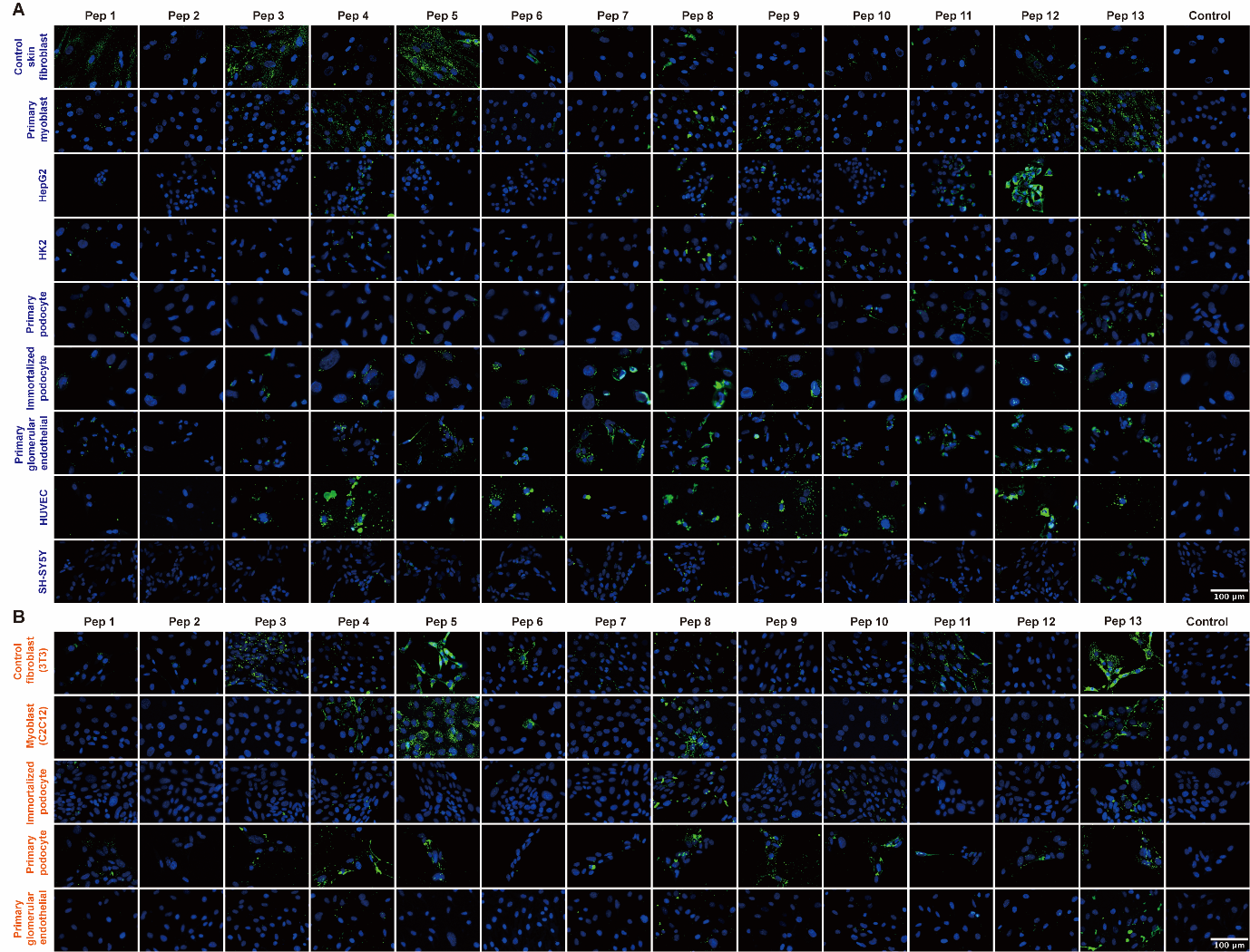

**Supplementary Figure S4: Cellular uptake of peptides across different human and mouse cell lines after 3 hours.**

(A) Representative fluorescence images showing uptake of FAM-labelled peptides in human cell lines, including control skin fibroblasts, primary myoblasts, HepG2, HK2, primary podocytes, immortalized podocytes, primary glomerular endothelial cells, HUVEC and SH-SY5Y cells. Nuclei were stained with DAPI (blue). Scale bars, 100 μm. (B) Representative fluorescence images showing uptake of FAM-labelled peptides in mouse cell types, including control fibroblasts (3T3), myoblasts (C2C12), immortalized podocytes, primary podocytes and primary glomerular endothelial cells. Nuclei were stained with DAPI (blue). Scale bars, 100 μm.

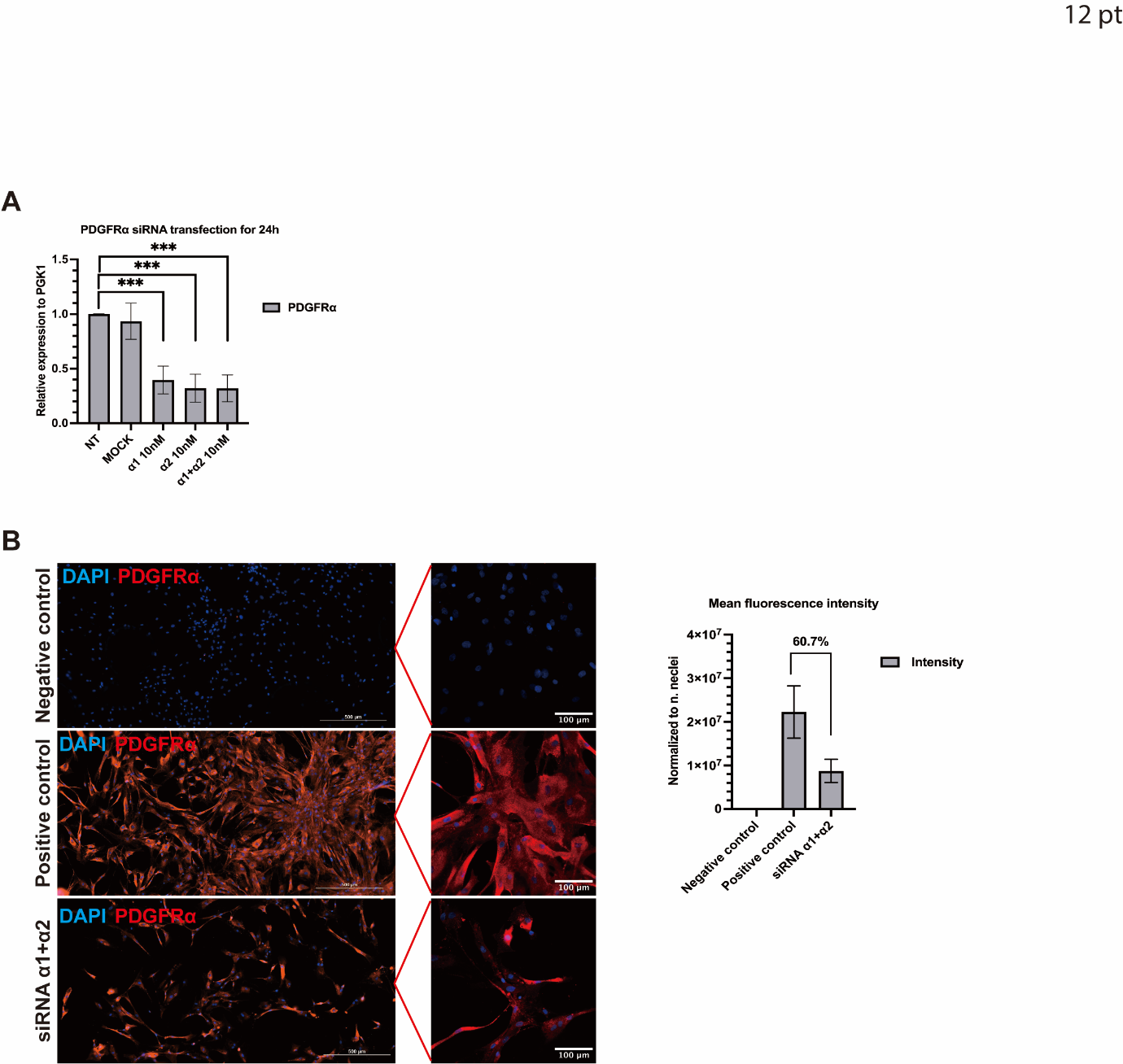

**Supplementary Figure S5: Validation of PDGFRα knockdown at mRNA and protein levels.**

(A) Relative PDGFRα mRNA expression following siRNA transfection (24 h) at indicated concentrations. Data are presented as mean ± SD. Statistical significance was determined using one-way ANOVA and post-Bonferroni test (***p< 0.001). (B) Immunofluorescence staining of PDGFRα (red) and nuclei (DAPI, blue) in control and siRNA-treated fibroblasts. Scale bars, 100 μm. Quantification of PDGFRα fluorescence intensity following siRNA treatment. Data are presented as mean ± SD.

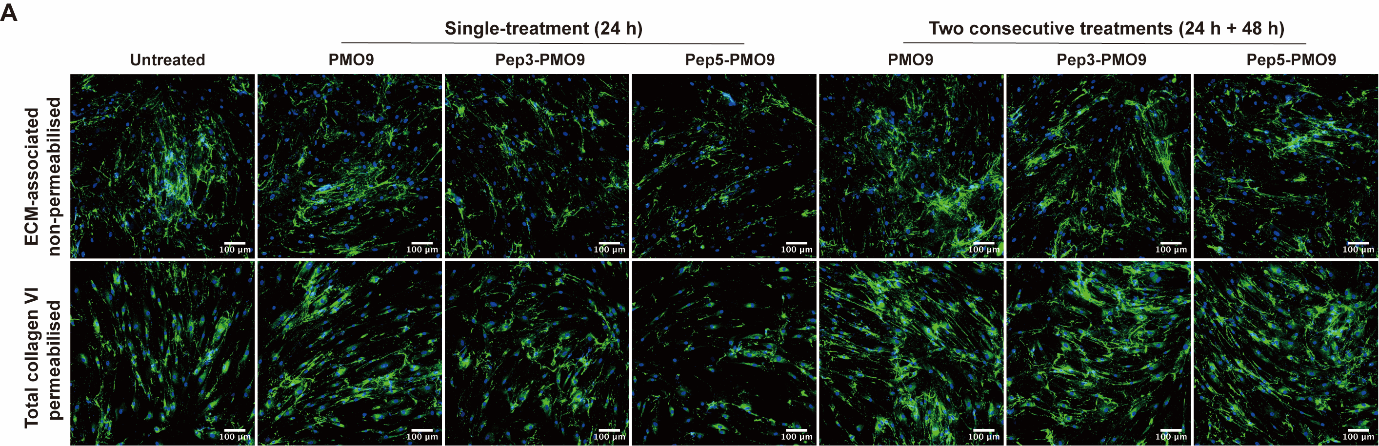

**Supplementary Figure S6: COL6 protein restoration after unconjugated and peptide-PMO conjugates in patient’s fibroblasts:**

Representative immunofluorescence images of collagen VI deposition following treatment with peptide–PMO conjugates in an additional patient fibroblasts.
Representative 20× immunofluorescence images showing ECM-associated collagen VI under non-permeabilised conditions and total collagen VI under permeabilised conditions following single treatment (24 h) or two consecutive treatments (24 h + 48 h) with unconjugated PMO9, Pep3-PMO9 or Pep5-PMO9 in patient 1 fibroblasts. Collagen VI staining is shown in green and nuclei were counterstained with DAPI (blue). Scale bar, 100 μm.

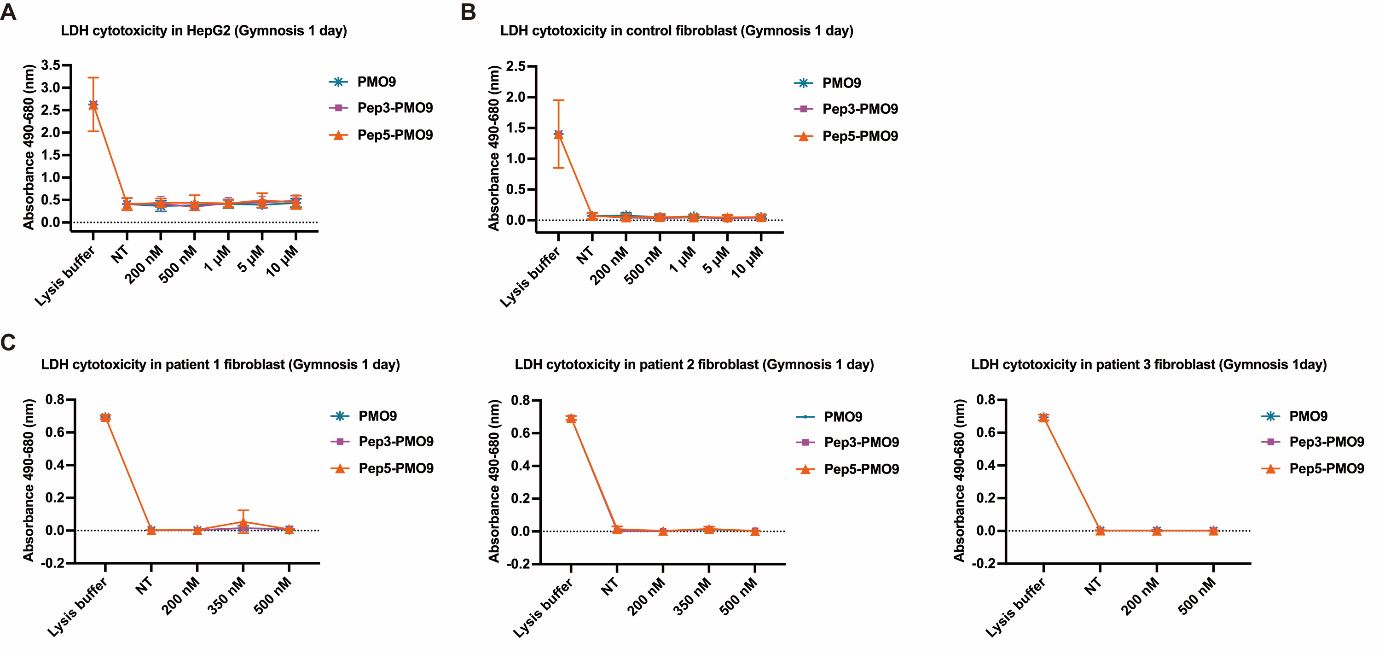

**Supplementary Figure S7: Cytotoxicity assessment of PMO and peptide-PMO conjugates.**

LDH cytotoxicity assay in HepG2 cells (A) and control fibroblasts (B) following gymnotic delivery for 24 hours with PMO9, Pep3-PMO9 or Pep5-PMO9 at the indicated concentrations. Lysis buffer was used as a positive control and NT indicate untreated group. (C) LDH cytotoxicity assay in fibroblasts derived from three COL6-CMD patients following gymnotic delivery for 24 hours with PMO9, Pep3-PMO9 or Pep5-PMO9 at the indicated concentrations.

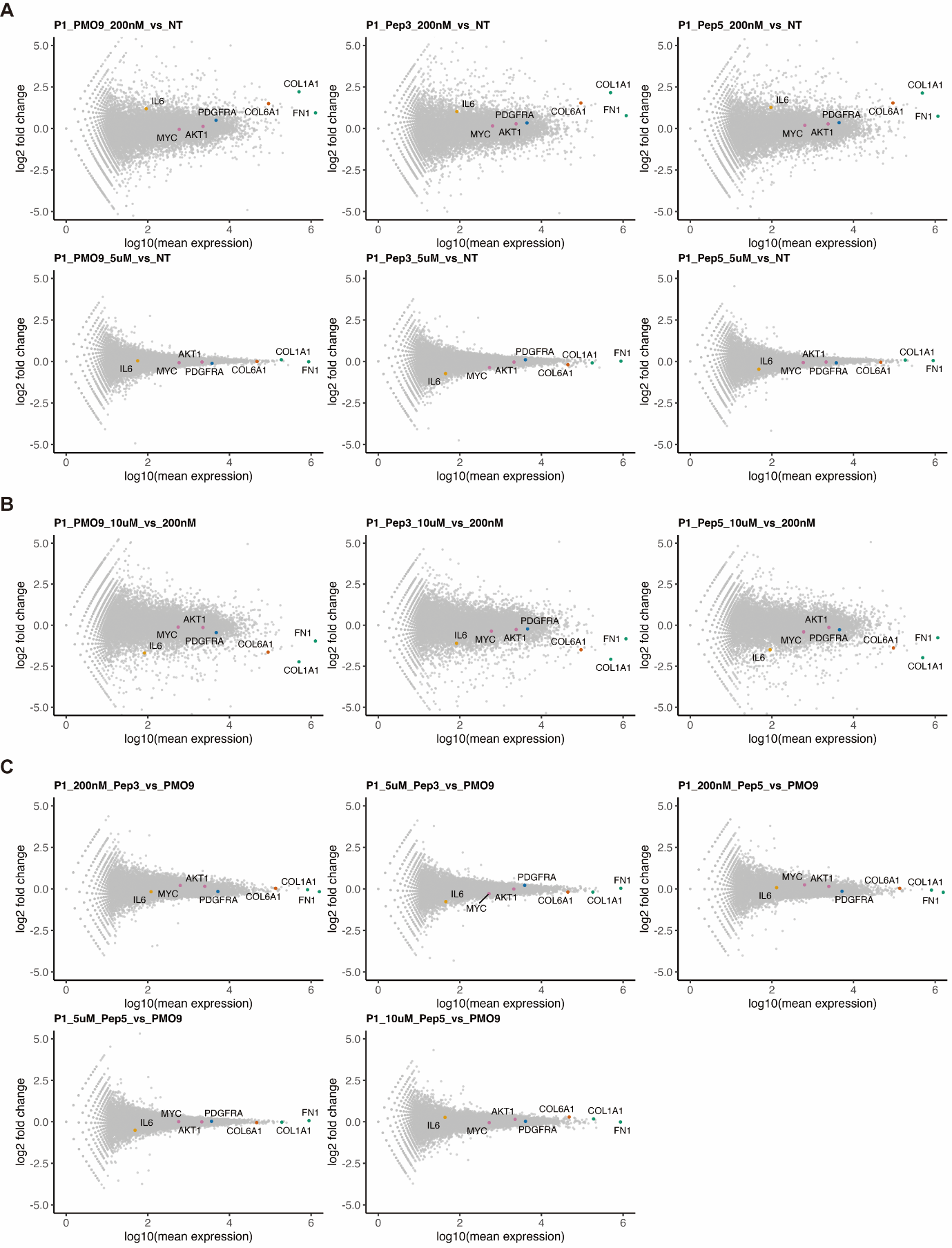

**Supplementary Figure S8: MA plots across treatment conditions in patient 1.**

(A–C) MA plots showing gene expression changes in patient 1 for selected comparisons. Each point represents a gene, plotted as log₂ fold change versus log₁₀ mean expression. Selected genes (*PDGFRA*, *AKT1*, *IL6*, *COL1A1*, *FN1*, *MYC*) are highlighted.

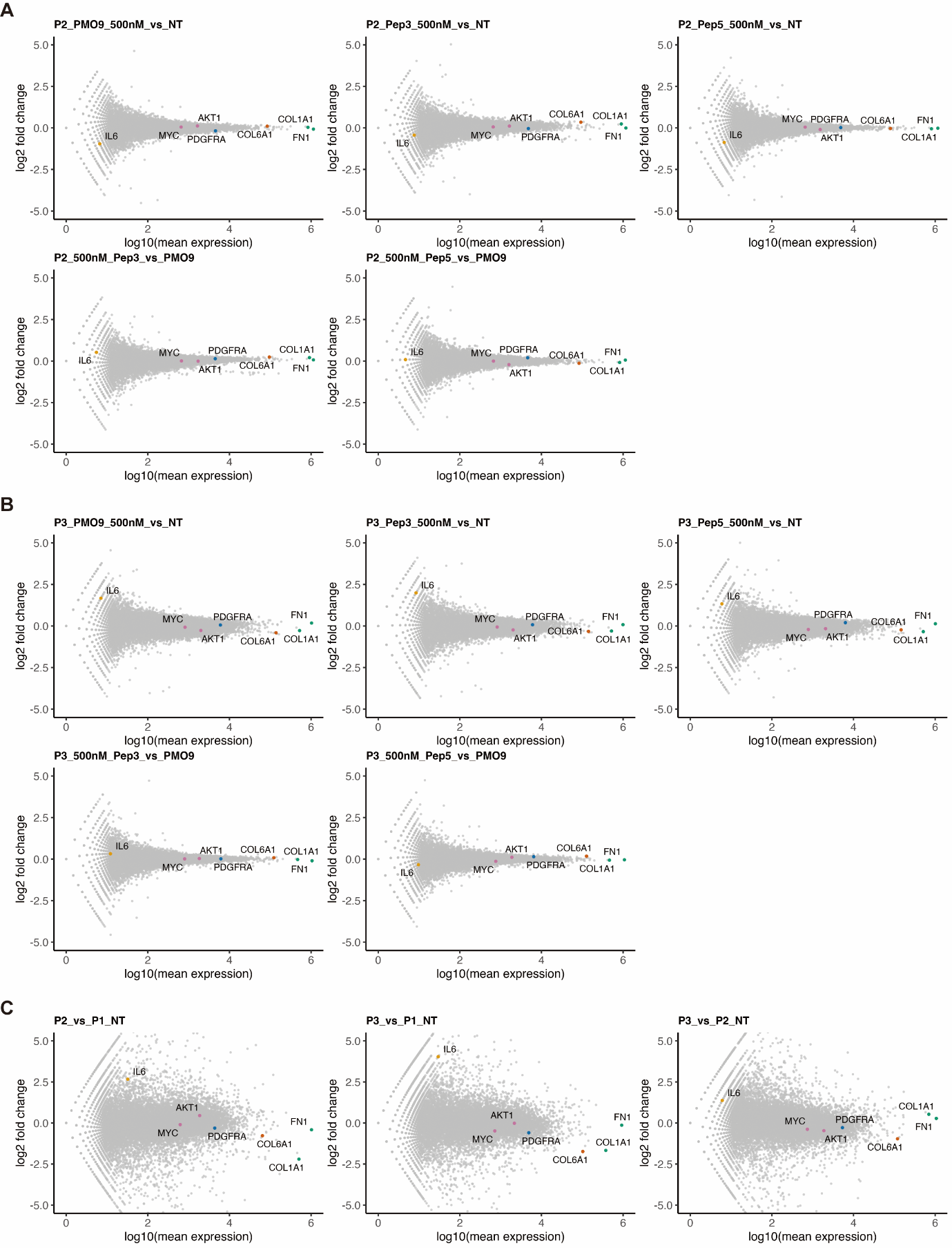

**Supplementary Figure S9: MA plots across treatment conditions in patients 2 and 3.**

(A–B) MA plots showing gene expression changes across indicated comparisons in patients 2 and 3. (C) MA plots comparing baseline gene expression between patient-derived fibroblast lines. Each point represents a gene, plotted as log₂ fold change versus log₁₀ mean expression. Selected genes are highlighted.

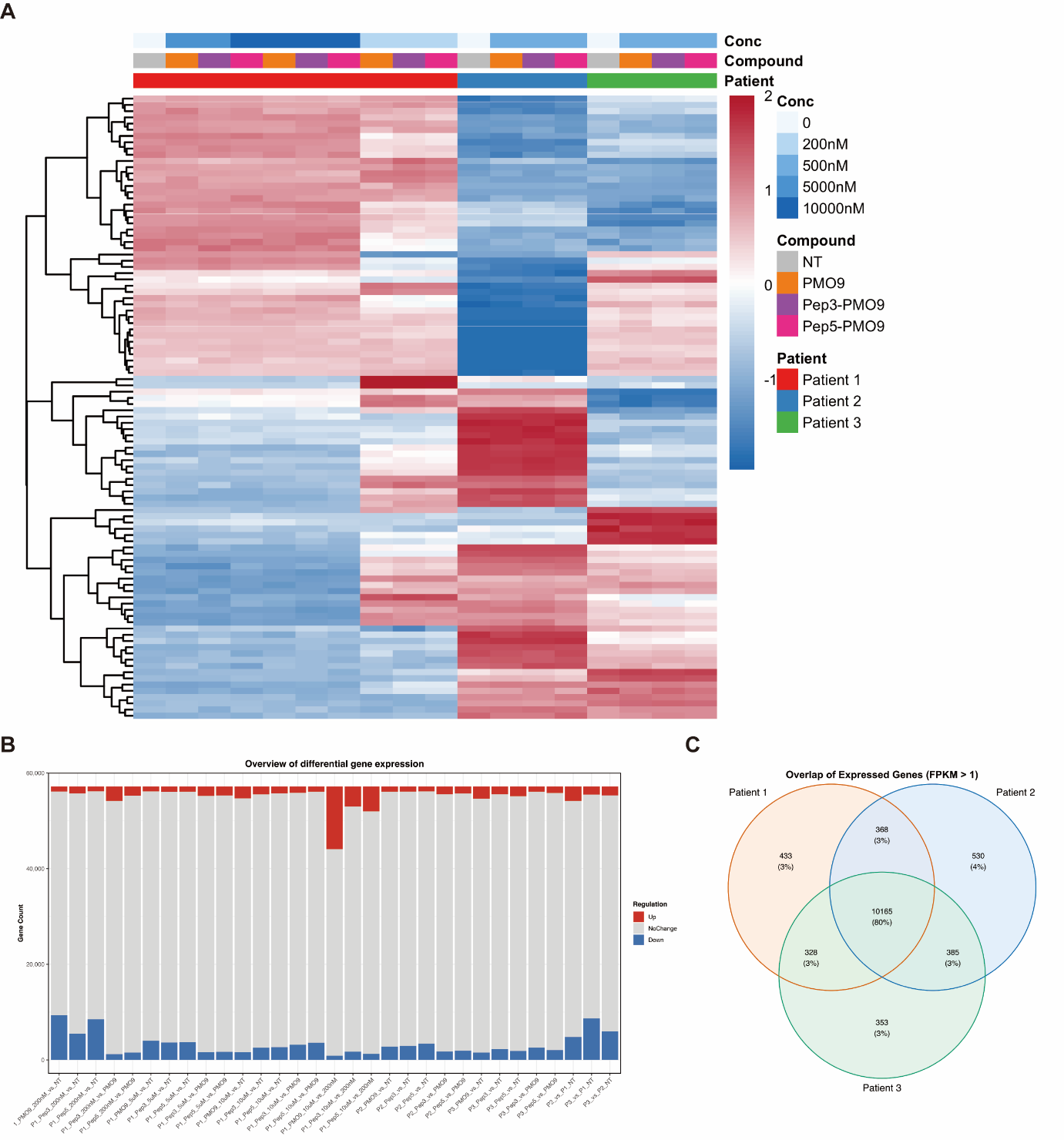

**Supplementary Figure S10: Extended overview of transcriptomic variation across samples.**

(A) Heatmap of the top variable genes across all samples based on variance-stabilized (VST) expression values. Samples are annotated by concentration, compound, and patient origin. (B) Overview of differential gene expression across all comparisons, showing the number of upregulated, downregulated, and unchanged genes. (C) Venn diagram showing overlap of expressed genes (FPKM > 1) across the three patient-derived fibroblast lines.

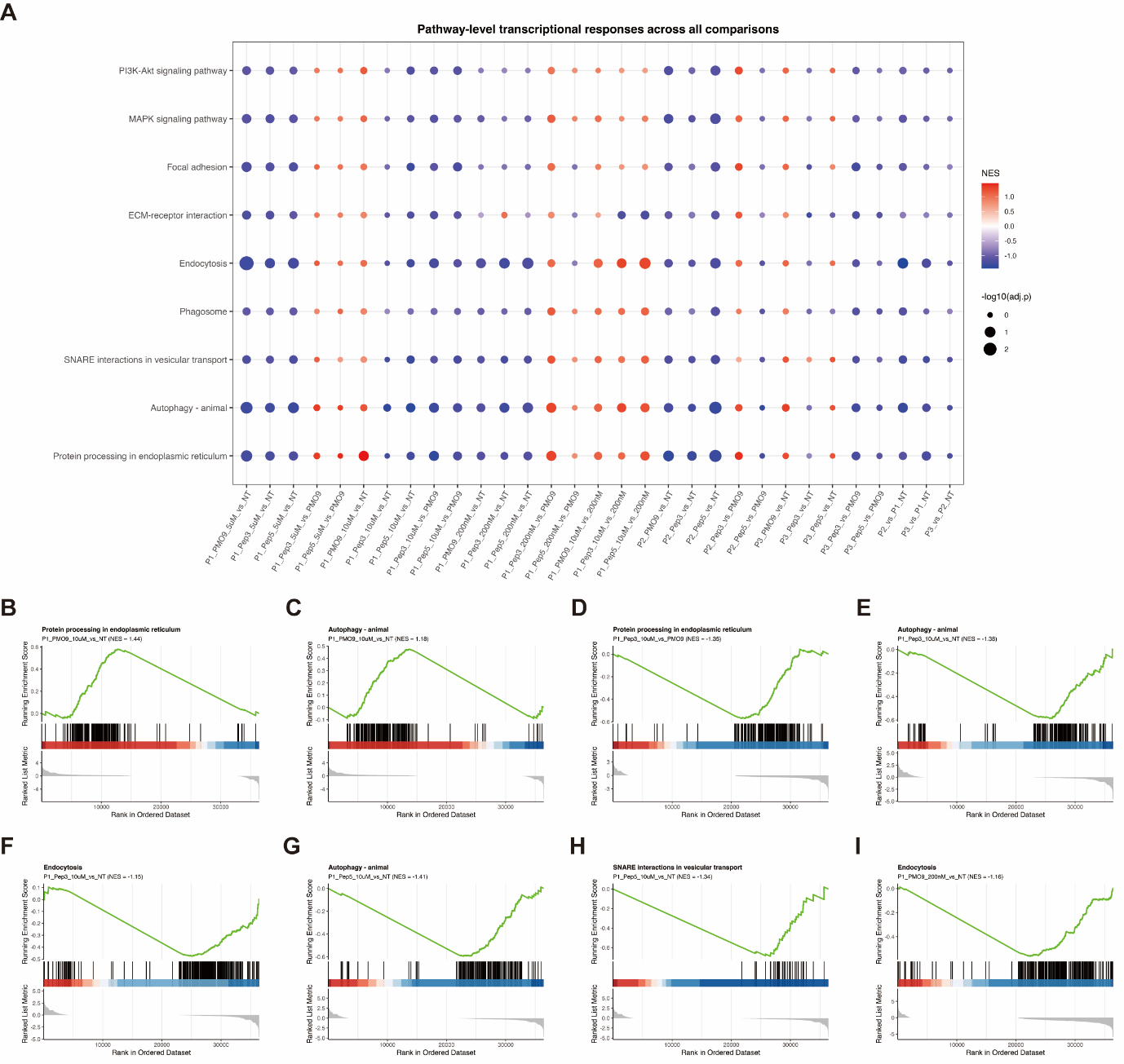

**Supplementary Figure S11: Pathway-level transcriptional responses across all comparisons**.

(A) Gene set enrichment analysis (GSEA) showing pathway-level transcriptional responses across all comparisons. Dot size represents –log₁₀(adjusted P value), and colour indicates normalized enrichment score (NES). (B–I) Representative enrichment plots for selected pathways across indicated comparisons. The running enrichment score is shown together with the distribution of genes contributing to the enrichment signal.
